# Enteropathogenic *Escherichia coli* (EPEC) Infection Induces Diarrhea, Intestinal Damage, Metabolic Alterations and Increased Intestinal Permeability in a Murine Model

**DOI:** 10.1101/2020.06.12.148593

**Authors:** Solanka E. Ledwaba, Deiziane V.S. Costa, David T. Bolick, Natasa Giallourou, Pedro H.Q.S. Medeiros, Jonathan R. Swann J, Afsatou N. Traore-Hoffman, Natasha Potgieter, James P. Nataro, Richard L. Guerrant

## Abstract

Enteropathogenic *E. coli* (EPEC) are recognized as one of the leading bacterial causes of infantile diarrhea worldwide. Weaned C57BL/6 mice pretreated with antibiotics were challenged orally with wild-type EPEC or *escN* mutant (lacking type 3 secretion system) to determine colonization, inflammatory responses and clinical outcomes during infection. Antibiotic disruption of intestinal microbiota enabled efficient colonization by wild-type EPEC resulting in growth impairment and diarrhea. Increase in inflammatory biomarkers, chemokines, cellular recruitment and pro-inflammatory cytokines were observed in intestinal tissues. Metabolomic changes were also observed in EPEC infected mice with changes in TCA cycle intermediates, increased creatine excretion and shifts in gut microbial metabolite levels. In addition, by 7 days after infection, although weights were recovering, EPEC-infected mice had increased intestinal permeability and decreased colonic claudin-1 levels. The *escN* mutant colonized the mice, but had no weight changes or increased inflammatory biomarkers, showing the importance of the T3SS in EPEC virulence in this model. In conclusion, a murine infection model treated with antibiotics has been developed to mimic many clinical outcomes seen in children with EPEC infection and to examine potential roles of selected virulence traits. This model can help in further understanding mechanisms involved in the pathogenesis of EPEC infections and potential outcomes and thus assist in the development of potential preventive or therapeutic interventions.

## INTRODUCTION

Gastroenteritis remains a major cause of morbidity and mortality in young children especially in developing countries [1]. Enteropathogenic *E. coli* (EPEC) has been recognized by the GEMS and MAL-ED studies as one of the major causes of moderate to severe diarrhea in children [2, 3]. Infection results in acute watery diarrhea accompanied by fever, vomiting and dehydration [4, 5].

EPEC contains the locus of enterocyte effacement regulator *(Ler)* gene, a major transcriptional activator of LEE open reading frames [6, 7]. EPEC virulence is mediated by the Type 3 secretion system (T3SS), characterized by attaching and effacing (A/E) lesions [6]. The T3SS consists of EPEC secreted components (Esc) and EPEC secretion proteins (Esp). In addition, *EscN* is the main driving force assisting in the ATPase response to enable activation of the T3SS, for efficient transportation of effector proteins into the enterocytes of the host [8]. During infection, EPEC attaches to epithelial cells via bundle forming pili *(bfp)* [9] followed by intimate adherence with the aid of the translocated intimin receptor *(tir)* and intimin (*eae*), which results in actin accumulation and formation of pedestal structures [5, 10]. EPEC is characterized by the presence (typical EPEC) or absence (atypical EPEC) of *bfp.* Typical EPEC are characterized by Localized Adherence (LA) *in vitro* [4] and have been reported to cause severe diarrhea in children under 12 months of age and in certain cases results in death [2, 3]. Atypical EPEC is characterized by LA-like [11], aggregative adherence or diffuse adherence *in vitro* [12, 13] and are increasingly being detected in children worldwide [14, 15].

Pathogens such as EPEC and EHEC compete with the resident microbiota for nutrients in order to colonize the intestinal environment. According to Freter’s nutrient niche, in order for microbes to be successful, it must have the capacity to grow fast in the intestine compared to its competitors [16]. These pathogens require the same carbon pathways which commensal *E. coli* uses, such as mannose and galactose *in vivo* [17].

EPEC have been studied extensively *in vitro*, which enables studies of localization traits, attaching and effacing lesions (A/E) and expression of the T3SS effector proteins [9, 18, 19]. *In vivo* studies have shown that a complete intestinal environment helps further determine the specific roles of EPEC traits involved in infections in animals and humans [20]. Animal models such as *Caenorhabditis elegans*, rabbits, pigs, and cattle have been used to study EPEC infections [21–24]. Infections induced by EPEC in C57BL/6 mouse models have also been reported [25–30], showing colonization of EPEC in the intestinal epithelial microvilli [25], changes in tight junction morphology and epithelial barrier function accompanied by inflammatory responses [27, 31]. These animal models have provided insights into the understanding of potential pathogenetic mechanisms of EPEC infection in humans. However, these models have not been able to replicate clinical outcomes observed in humans. We describe a murine model in which the microbiota have been disrupted via broad-spectrum antibiotic cocktail to enable efficient colonization and clinical outcomes of EPEC infection in mice promoting growth impairment, diarrhea, intestinal damage, metabolic alterations and increased intestinal permeability.

## RESULTS

### EPEC infection leads to growth impairment and diarrhea

A murine EPEC infection model able to induce changes in body weight and diarrhea, which are important outcomes in children infected by EPEC, has been needed [20]. Depletion of intestinal microbiota by antibiotics has been shown to be effective in promoting colonization by bacterial pathogens [32–34]. We therefore, tested whether pretreatment with antibiotics could enable the study of body weight and diarrhea in mice infected with EPEC (10^10^ CFU) (Fig 1A). EPEC inhibited the growth of mice when compared to the control group from days 2-5 post infection (p.i.) (p<0.05, Fig 1B). EPEC infection also induced moderate to severe diarrhea at day 3 p.i. (p<0.0001), persisting until day 5 p.i. (Fig 1C). The changes in body weight exhibited by EPEC-infected mice were correlated with diarrhea scores (Fig 1D), showing higher diarrhea scores were associated with greater weight shortfalls.

**Fig 1.**
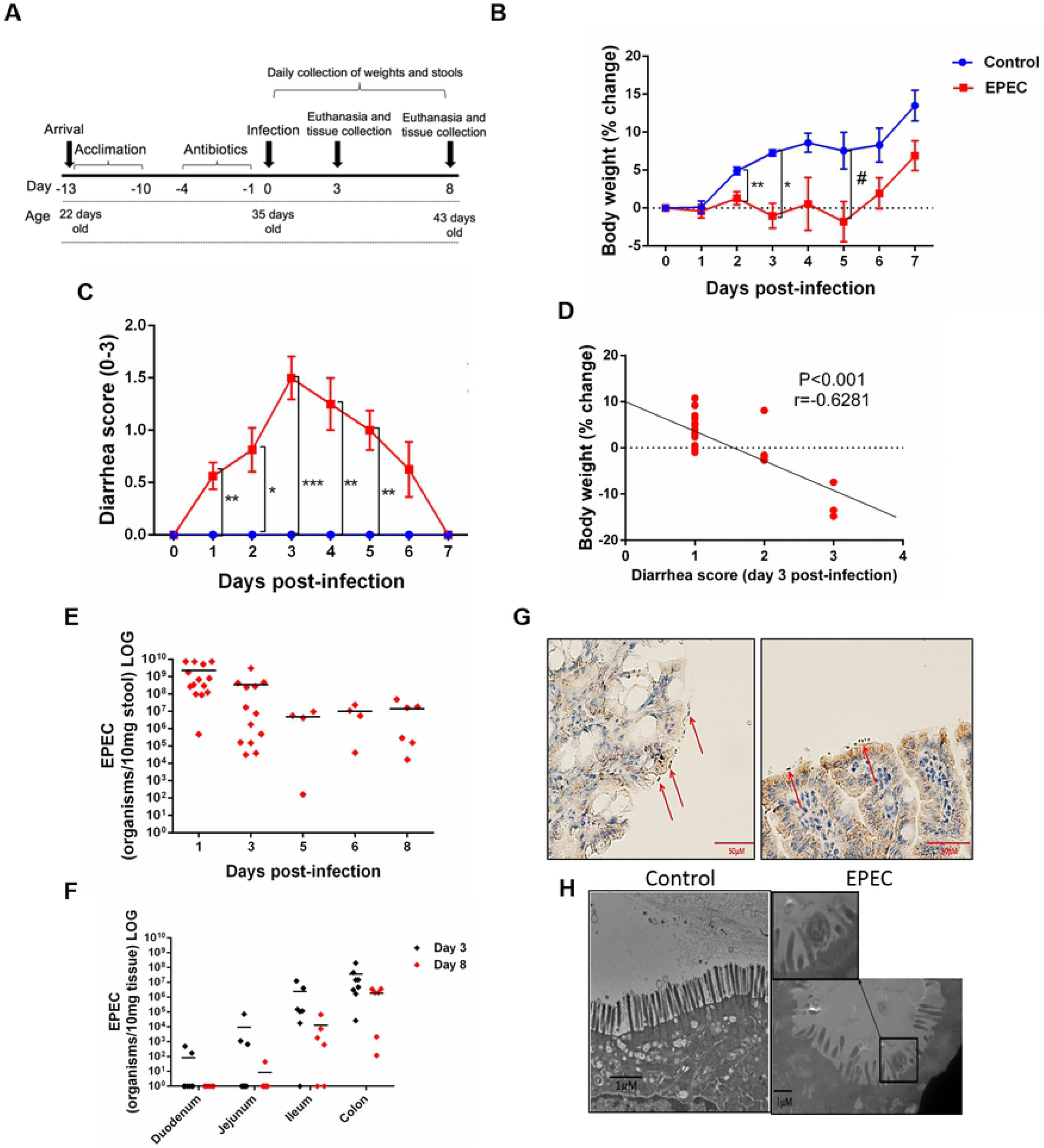
EPEC infection impairs weight gain, induces diarrhea and colon colonization in C57BL/6 mice. (**A**) Experimental timeline of EPEC infection model. Weaned C57BL/6 mice pretreated with antibiotic cocktail were orally infected with EPEC (10^10^ CFU). (**B**) Change in body weight of C57BL/6 mice infected with EPEC (EPEC) or uninfected (Control) (n=12/group). Line graphs represents mean±SEM. **p < 0.009, *p<0.001 and #p < 0.04 using multiple Student’s *t-test.* (**C**) Diarrhea score of EPEC and control mice. Line graphs represent median±SEM. *p<0.01, **p<0.006 and ***p<0.0001 using Kruskal-Wallis test followed by Dunn’s test. (**D**) Significant negative correlation between diarrhea score and change in body weight at day 3 p.i. (Spearman rank test). (**E**) Quantification of EPEC shedding in stools. (**F**) Quantification of EPEC tissue burden in the intestinal tissues (duodenum, jejunum, ileum and colon) at day 3 and 8 p.i. (**G**) Immunohistochemistry staining of intimin (red arrows), on ileal tissue of EPEC infected mice at day 3 p.i. (**H**) Transmission electron microscope (TEM) at day 3 p.i., showing disruption of the microvilli of mice infected with EPEC.

### EPEC colonizes the ileum and colon in mice

In order to confirm that growth impairment and diarrhea were promoted by EPEC infection, DNA was extracted from stools of EPEC-infected mice and quantitative PCR was used to measure shedding. As shown in Fig 1E, most of the EPEC-infected mice exhibited 10^8^-10^10^ organisms/10 mg stool at day 1 p.i. and less shedding was observed at day 8 p.i.

Given that EPEC is an intestinal pathogen, we further investigated which intestinal sections were predominantly colonized by EPEC using quantitative PCR to measure tissue burdens. EPEC was found to predominantly colonize the ileum (<10^7^ organisms/mg tissue) and colon (<10^8^ organisms/mg tissue) of mice at day 3 pi (Fig 1F), and the same trend was also observed at day 8 p.i. Fig 1G shows intimin-stained EPEC adherence on blunted ileal mucosa, with disruption of the microvilli shown by TEM (Fig 1H).

**Fig 2.**
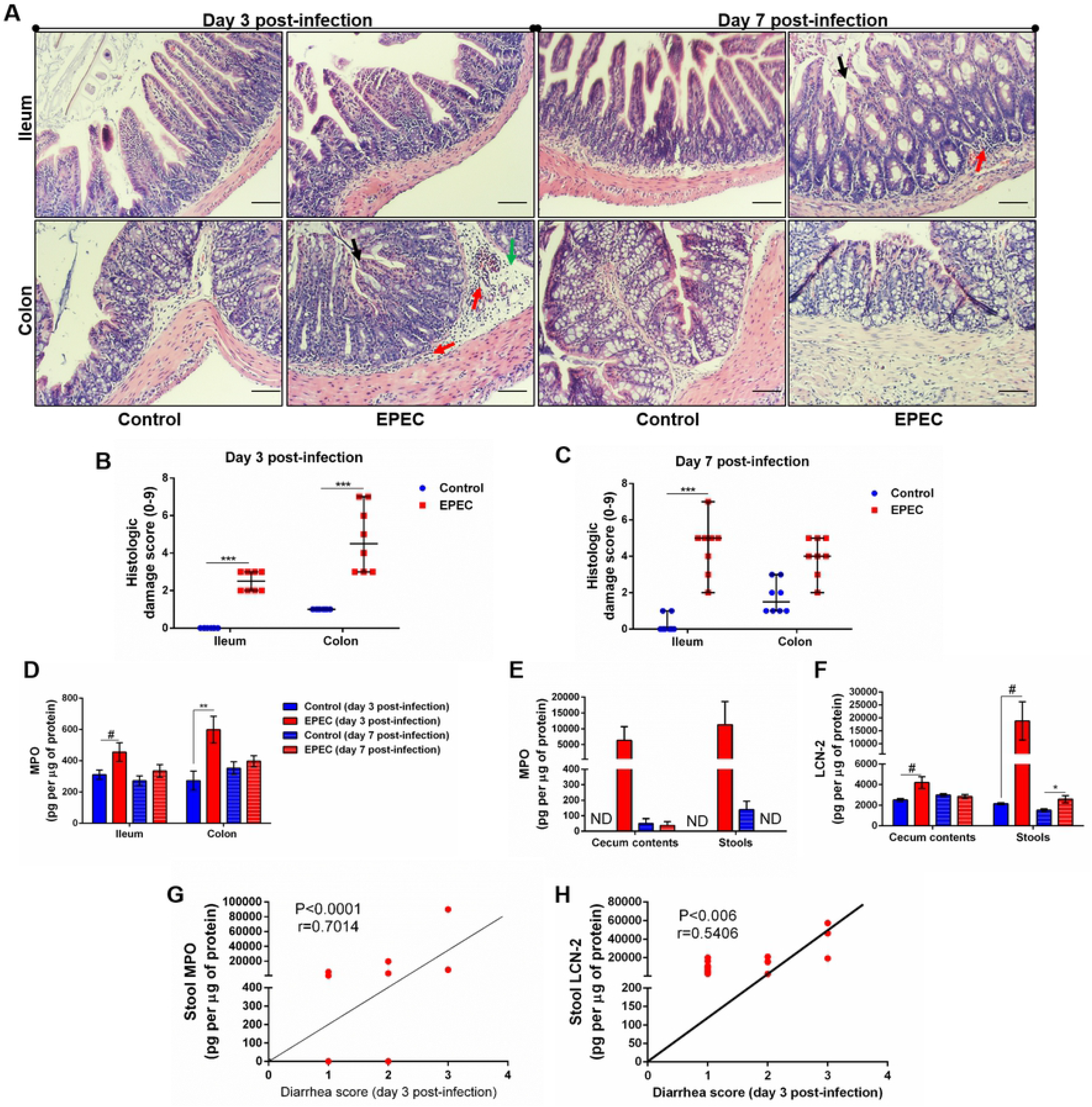
EPEC infection model induces acute colonic damage and inflammation followed by a positive correlation between diarrhea and inflammation markers in stools. (**A**) Representative H&E stains of ileal and colonic tissue collected from control and EPEC infected mice at day 3 and 7 p.i. Scale bars, 100μm. (**B-C**) Histologic damage score based on epithelial damage (black arrow), mucosal edema (green arrow) and inflammatory cell infiltrate (red arrow) in the ileal and colonic tissue of uninfected (control group) and EPEC-infected (EPEC group) (**B**) mice at day 3 and (**C**). mice at day 7 p.i. Bars represent median±SEM (n=8). ***p<0.0001 using Kruskal-Wallis test followed by Dunn’s test. (**D**) MPO levels measured in intestinal tissue lysates (ileum and colon) collected at day 3 and day 7 p.i. (**E**) MPO and (**F**) LCN-2 levels measured in the cecal and stool lysates collected at day 3 and day 7 p.i. Bars represent mean±SEM (n=8). # p<0.05, **p<0.03 and p<0.007 using multiple Student’s *t-test.* (**G**) Significant positive correlation between diarrhea score and MPO levels in stools of EPEC-infected mice at day 3 p.i. (Spearman rank test). (**H**) Significant positive correlation between diarrhea score and lipocalin levels in stools of EPEC-infected mice at day 3 p.i. (Spearman rank test).

These findings indicate that EPEC infection promotes a self-limited symptomatic acute disease in antibiotic-pretreated mice. Fecal shedding of EPEC, and tissue burdens were detected to day 8 p.i. in infected mice.

### EPEC infection promotes acute intestinal tissue damage and inflammation

Given that our EPEC infection model resulted in a significant colonization in the ileum and colon, we next investigated whether EPEC infection promotes ileal and colonic damage. The ileal and colonic histological damage in EPEC-infected mice was characterized by the loss of epithelial integrity, moderate edema in the submucosa and infiltration of inflammatory cells into the lamina propria and submucosa, with significant histology score differences from controls in both ileum and colon at day 3, and persistent significant differences in the ileum extending to day 7. The damage was found to be greater in colon compared to control mice (Fig 2A) at day 3 p.i., as confirmed by measurement of histologic damage score (p<0.0001, Fig 2B). On day 7 p.i., the damage in the ileum of EPEC-infected mice was higher when compared to the control mice (p<0.0001, Fig 2A and 2C).

Myeloperoxidase (MPO), a marker of neutrophil activity in intestinal mucosa, and lipocalin-2 (LCN-2), a glycoprotein upregulated in tissue damage under infection conditions, have been considered as biomarkers of environmental enteric dysfunction, including EPEC, in children [35–38]. To ensure that our EPEC infection model mimics the alterations of these biomarkers as observed in children, we measured MPO and LCN-2 in the ileal and colonic tissues, as well as in cecal contents and stools. We found increased MPO levels in ileum and colon tissues of EPEC-infected mice at day 3 p.i. when compared to the control group (p<0.05 and p<0.03 respectively, Fig 2D). Of note, a trend of increase in MPO levels was also observed in the cecal contents and stools at day 3p.i. (Fig 2E), however no statistical significance was found. On day 7 p.i., MPO levels were reduced in the intestinal (ileum and colon), cecal contents and stools of EPEC-infected mice compared to controls (Fig 2D-E). However, increased LCN-2 levels were found in cecal contents (day 3 p.i.) and stools (day 3 and 7 p.i.) of EPEC-infected mice when compared to control mice (p<0.05-day 3 p.i. or p<0.03-day 7 p.i., Fig 3F).

**Fig 3.**
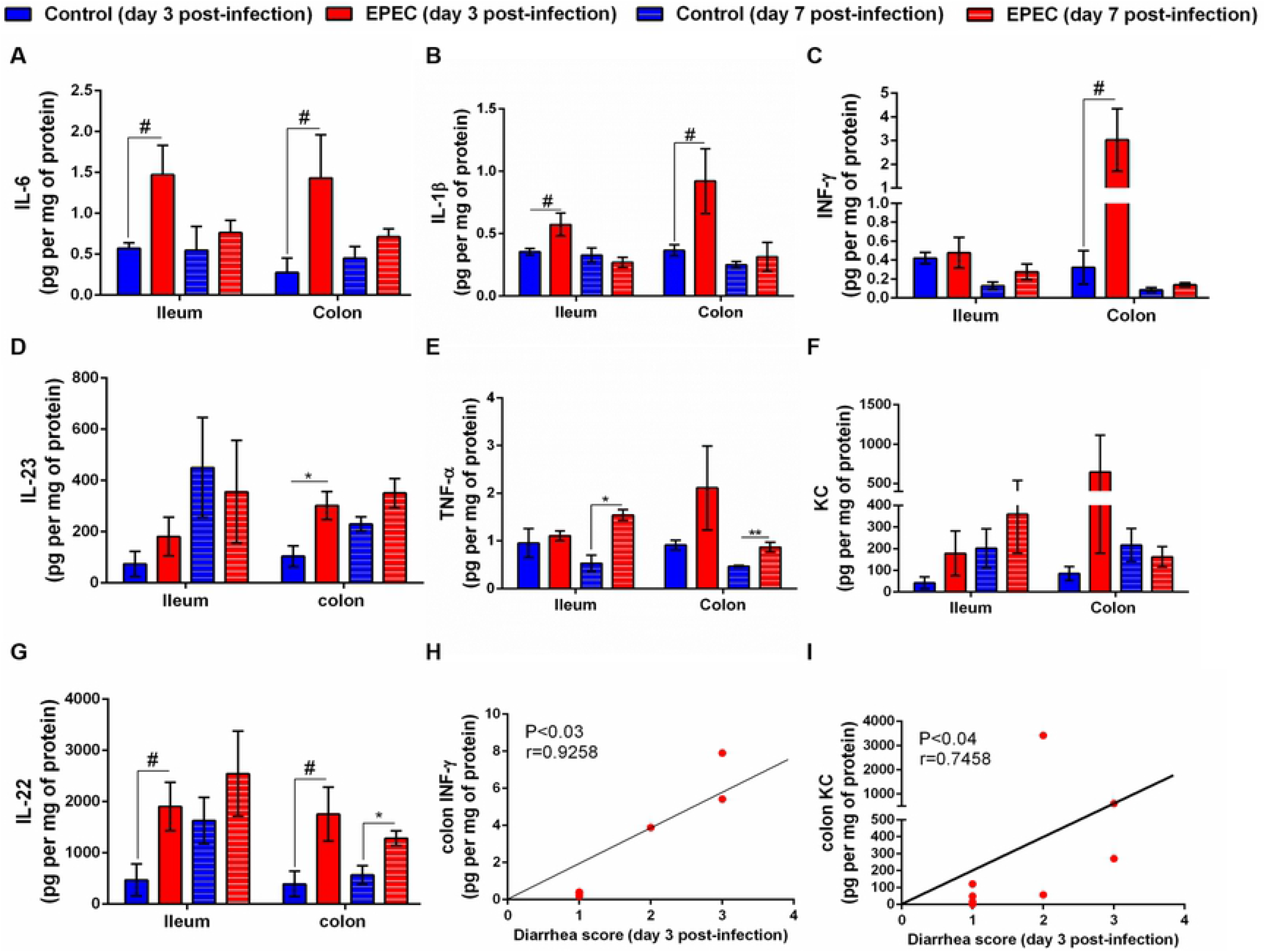
EPEC infection increases pro-inflammatory mediators, and diarrhea scores correlates positively with INF-γ and KC levels in colonic tissues of EPEC infected mice. (**A**) Levels of IL-6, (**B**) IL-1β, (**C**) INF-γ, (**D**) IL-23, (**E**) TNF-α, (**F**) KC and (**G**) IL-22 in the ileal and colonic tissues of control and EPEC infected mice at day 3 and day 7 p.i. were measured by ELISA. Bars represent mean±SEM (n=8). # p<0.05, *p<0.01 and **p<0.001 using multiple Student’s *t-test.* (**H**) Significant positive correlation between diarrhea score and INF-γ levels in colonic tissues of EPEC infected mice at day 3 p.i. (Spearman rank test). (**I**) Significant positive correlation between diarrhea scores and KC levels in colonic tissues of EPEC-infected mice at day 3 p.i. (Spearman rank test).

We correlated diarrhea score with MPO or LCN-2 levels in stools samples of EPEC-infected mice at day 3 p.i., when mice exhibited higher diarrhea score, a positive correlation was found between diarrhea score and MPO levels in stools (p<0.0001, r=0.7014, Fig 2G). Positive correlation was also observed on diarrhea score and LCN-2 levels (p<0.006, r=0.5406, Fig 2H). These data indicated that high diarrhea scores are associated with increased MPO and LCN-2 levels.

### EPEC infection alters pro-inflammatory and anti-inflammatory cytokine synthesis in ileum and colon in a stage disease-dependent manner in mice

Next, we further analyzed which pro-inflammatory (IL-6, IL-1β, INF-γ, IL-23, tumor necrosis factor-α (TNF-α), KC-keratin chemoattractant-, IL-17, IL-18) and anti-inflammatory (IL-22, IL-33, IL-10) cytokines were affected by EPEC infection during acute (day 3 p.i.) and later (day 7 p.i.) stage of the disease. As shown in Fig 3, higher levels of IL-6 (p<0.05), IL-1β (p<0.05), INF-γ (p<0.05), IL-23 (p<0.01) and IL-22 (p<0.05) were detected in colonic tissues of EPEC-infected mice and were significantly different when compared to controls (Fig 3A-D and 3G). EPEC infection in the ileal tissue resulted in significant increase in IL-6, IL-1β and IL-22 when compared to control mice (p<0.05, Fig 3A, 3B and 3G). In the later stage of disease (day 7 p.i.), TNF-α (ileal and colonic tissues) and IL-22 (colonic tissues) were significantly increased in EPEC-infected mice when compared to control mice (p<0.01, Fig 3E and 3G). In addition, increase of KC in the ileal and colonic tissues was observed, however, not statistically significant (Fig 3F).

We analyzed the correlation of diarrhea severity with the levels of INF-γ or KC in colonic tissues of EPEC-infected mice and found a strong positive correlation between diarrhea score and colonic INF-γ levels (p<0.03, r=0.9258, Fig 3H) or KC levels (p<0.04, r=0.7458, Fig 3I). We did not find any change in the levels of IL-17, IL-18, granulocyte-macrophage colony-stimulating factor (GMCSF), IL-33 and IL-10 in ileal and colonic tissues samples of EPEC-infected mice compared to the controls (Supplementary Fig1A-E).

Taken together, these data indicate that EPEC infection promotes two-phase of disease in mice: an acute stage characterized by growth impairment, intestinal damage, presence of diarrhea and increased of MPO, LCN-2, IL-6, IL-1β, INFγ, IL-23 and IL-22; and a later stage with an increase in LCN-2 and TNF-α levels in an absence of diarrhea. In addition, the data suggests that the colon is more affected by EPEC infection in mice.

### EPEC-infected mice with diarrhea demonstrates upregulation of pro-inflammatory cytokines, inflammatory markers, STAT and apoptosis markers in colon

Due to an increase in diarrhea severity and colonic INF-γ levels that were positively correlated on day 3 p.i., we evaluated the profile of gene expression from the colon tissues of EPEC-infected mice with diarrhea and controls using Taqman assay. In total, 92 genes were evaluated, among these 37 were upregulated and four were downregulated (p<0.05, Fig 4). INF-γ, GZMB, CXCL10, IL-6, IL-1β were the most upregulated genes, showing approximately 85, 30, 27, 23 and 16-fold-change in relation to controls (p<0.05, Fig 5A). In mice with diarrhea, EPEC infection resulted in upregulation of pro-inflammatory mediators *(INF-γ, TNF-α, TNFRS, IL-1α, IL-1β, IL-2Rα, IL-5, IL-6 and IL-12b*, Fig 4B), chemokines *(CCL2, CCL5, CCL19, CXCL10* and *CXCL11*, Fig 4C), chemokine receptors *(CCR2* and *CCR7*, Fig 4C), cellular recruitment *(VCAM1, ICAM1* and *SELP*, Fig D), *CD68* (a macrophage marker, Fig 4E), *CD8ε* (marker of T-cell activation, Fig 4E), inflammation markers *(C3, CD38, CD40, CSF2, GZMB* and *MD2*, Fig 4F), transcription factors (phosphorylated-signal transducer and activator of transcription-1-STAT1-, STAT3 and STAT4, Fig 4G) and apoptosis markers (Fas and Bax, Fig 4I) in colon of mice on day 3 p.i. In addition, EPEC-infection also upregulated gene expression of anti-inflammatory mediators, such as TGF-β1, HMOX1, PTPRC, SOCS1 and LIF when compared to control mice (p<0.05, Fig 4J).

**Fig 4.**
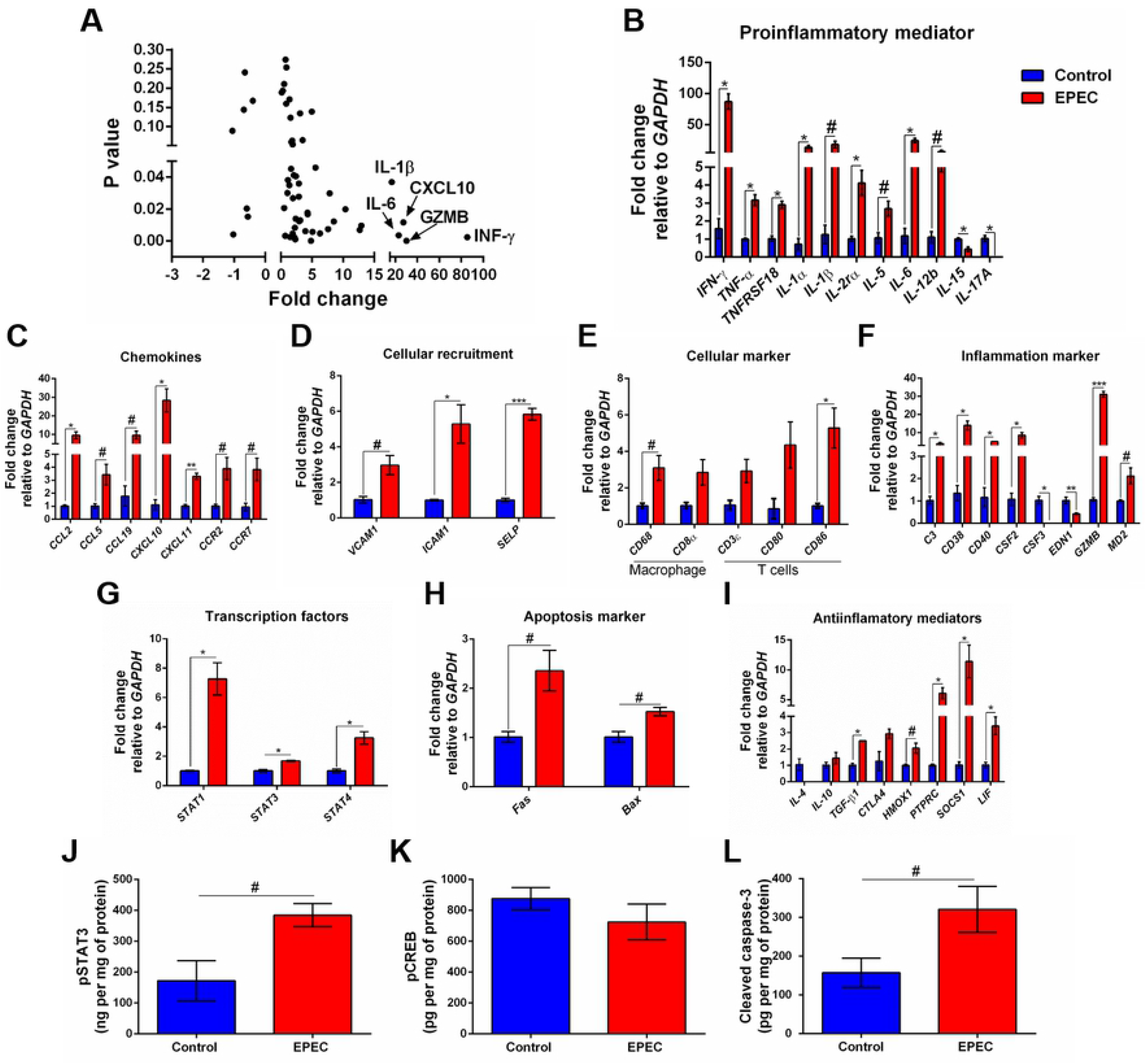
EPEC-infected mice with soft or unformed stools exhibit higher expression of mRNA for pro-inflammatory mediators, chemokines and other inflammatory markers. (**A**) mRNA expression fold changes (on the x axis) and the corresponding P values (on the y axis). Gene names are labeled next to highlighted significant results. Taqman qPCR analysis of (**B**) pro-inflammatory, (**C**) chemokines, (**D**) cellular recruitment, (**E**) cellular marker, (**F**) inflammation marker, (**G**) transcription factors, (**H**) apoptosis marker and (**I**) anti-inflammatory genes in the colonic tissues of control and EPEC-infected mice with moderate (score=2) or severe (score=3) diarrhea scores at day 3 p.i. (**A-I**) Expression levels were normalized with GAPDH, as an internal housekeeping gene. Bars represent mean±SEM (n=3). # p<0.05, *p<0.01, **p<0.001 and ***p<0.0001 using multiple Student’s *t-test.* Levels of (**J**) pSTAT3, (**K**) pCREB and (**L**) cleaved caspase-3 in colonic tissue lysates of control and EPEC-infected mice at day 3 p.i. measured using ELISA. Bars represent mean±SEM (n=8). # p<0.05 using Student’s *t-test.*

STAT3 is a transcription factor involved in response of cytokines such as IL-6, and its activation results in expression of target genes involved in inflammatory and anti-inflammatory responses [39, 40]. To investigate the levels of phosphorylated STAT3 (pSTAT3), its active form, we found increased levels of pSTAT3 in the colon of mice infected with EPEC on day 3 p.i. (p<0.05, Fig 4J).

CREB is another transcription factor involved on the transcription of inflammatory (such as IL-6, IL-2 and TNF-α) and antiinflammatory (IL-4, IL-10 and IL-13) mediators [41, 42]. To determine the levels of phosphorylated cyclic AMP –responsive element-binding protein (pCREB), its active form, the difference between EPEC-infected mice and controls on day 3 p.i. was not found (Fig 4K).

Given that EPEC infection increased the gene expression of apoptosis markers, we further evaluated the levels of cleaved caspase-3 using ELISA. We found that EPEC infection induced an increase on cleaved caspase-3 in the colonic tissues when compared to control mice at day 3 p.i.(p<0.05, Fig 4L), confirming an apoptosis process during EPEC infection.

### EPEC infection model induces metabolic perturbations

Metabolic perturbations following EPEC infection were further analyzed using Orthogonal projection to latent structures discrimination analysis (OPLS-DA). Urinary metabolic profiles of each of the mice infected with EPEC were compared to the age-matched uninfected mice at days 1 and 3 p.i. No differences were observed between the controls and EPEC infected mice on the day 1 p.i. (Fig 5A).

**Fig 5.**
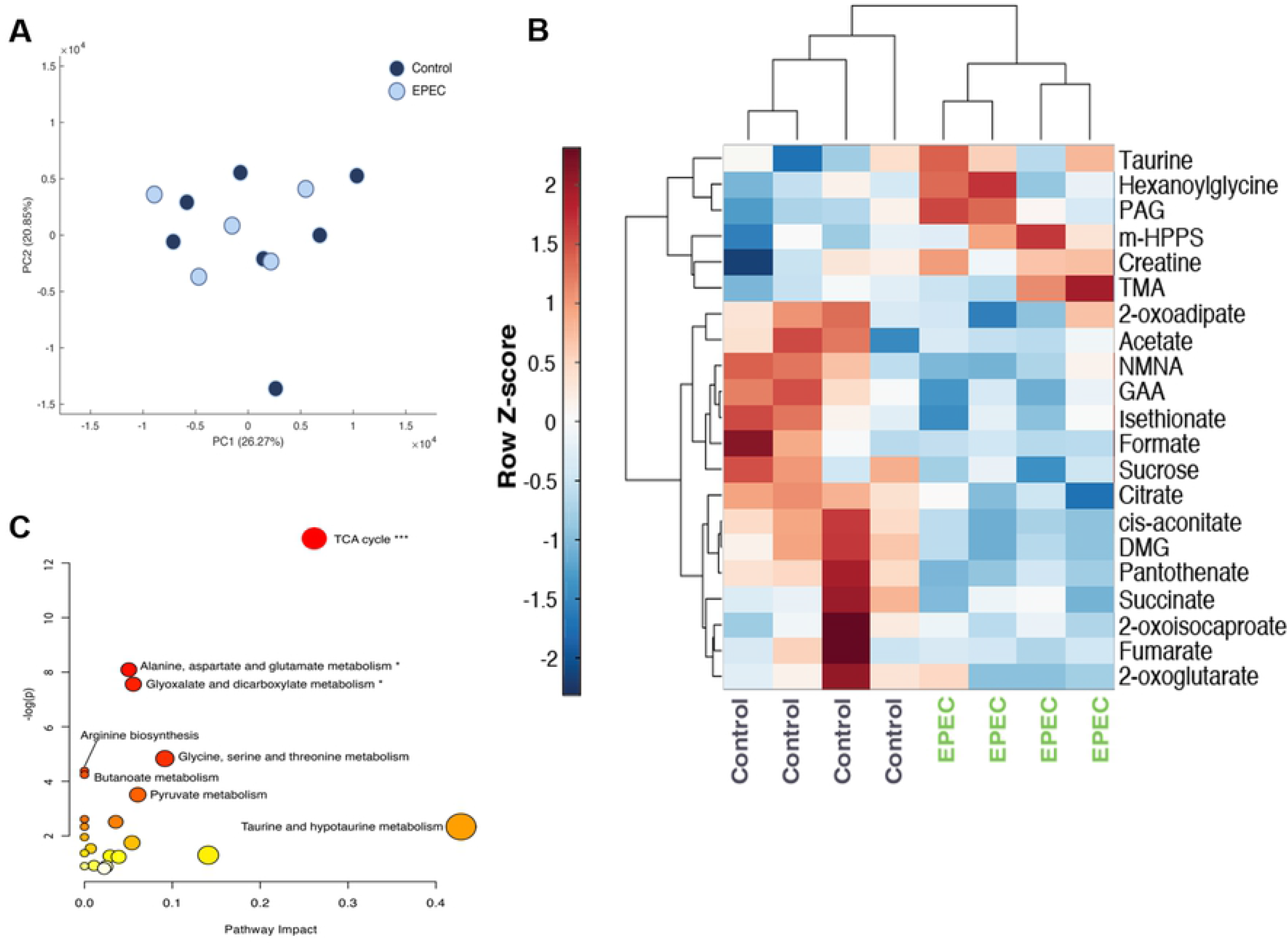
Metabolomic analysis of urinary specimens collected from age-matched, control and EPEC infected mice. (**A**) Principal component analysis (PCA) scores plot at day p.i. showing no difference between infected and uninfected mice (control n=7, EPEC n=5). (**B**) Unsupervised hierarchical clustering and heat-map of urinary metabolites from control and EPEC-infected mice at day 3 p.i. (n=4). Each row represents a metabolite and each column represents a mouse sample. The row Z-score (scaled expression value) of each metabolite is plotted in red-blue colour. The red colour indicates metabolites, which are high in abundance and blue indicates metabolites low in abundance. DMG, dimethylglycine; GAA, guanidinoacetic acid; m-HPPS, m-hydroxyphenylpropionylsulfate; NMNA, N-methyl-nicotinic acid; PAG, phenylacetylglycine; TMA, trimethylamine. (**C**) Pathway analysis plot created using MetaboAnalyst 4.0 showing metabolic pathway alterations induced by EPEC. Each pathway is represented as a circle. Darker colors indicate more significant changes of metabolites annotated on the corresponding pathway. The x-axis represents the pathway impact value computed from pathway topological analysis, and the y-axis is the-log of the p-value obtained from pathway enrichment analysis. The pathways that were most significantly changed are characterized by both a high-log(p) value and high impact value (top right region). FDR adjusted p-values, *p<0.05, ***p<0.0001.

EPEC infection-driven metabolic variation was observed at day 3 pi (OPLS-DA model: Q^2^Y = 0.57, P = 0.027 (1000 permutations). EPEC infection resulted in reduced excretion of TCA cycle metabolites (succinate, cis-aconitate, citrate, 2-oxoglutarate and fumarate) and choline related metabolites [dimethylglycine (DMG) and trimethylamine (TMA)] (Fig 5B). Lower urinary excretion of the tryptophan catabolite N-methylnicotinic acid (NMNA), the creatinine precursor guanidinoacetic acid (GAA) and the amino acid catabolites 2-oxoisocaproate, 2-oxoadipate were also observed. Isethionate, formate, pantothenate, and sucrose were also excreted in lower amounts by EPEC infected mice. Increases in the excretion of gut microbial-derived metabolites [acetate, phenylacetylglycine (PAG), m-hydroxyphenylpropionylsulfate (m-HPPS)], were observed in infected mice. Urinary excretion of taurine, creatine and b-oxidation product hexanoylglycine, were also elevated at day 3 p.i (Fig 5B). Pathway analysis revealed that the TCA cycle was the biochemical pathway most influenced by the infection (Impact: 0.26, p-value: 2.5E-6, FDR adjusted p-value: 2.1E-4) (Fig 5C).

### EPEC infection increases intestinal permeability and decreases colonic claudin-1 expression in mice

Alteration on intestinal permeability related to EPEC infection has been reported in children [43]. Given that the intestinal tissues of EPEC-infected mice showed increase in inflammatory markers on later stage of disease, we investigated whether intestinal permeability was altered by using a FITC dextran assay in our experimental model. As shown in Fig 6A, EPEC infection resulted in increased levels of plasma 4kDa FITC dextran, indicating higher intestinal permeability when compared to control mice at day 7p.i. (p<0.006). A strong positive correlation was found between the levels of plasma 4kDa FITC dextran and colonic INF-γ at day 7 p.i. in EPEC-infected mice (p<0.003, r=0.9219, Fig 6B).

**Fig 6.**
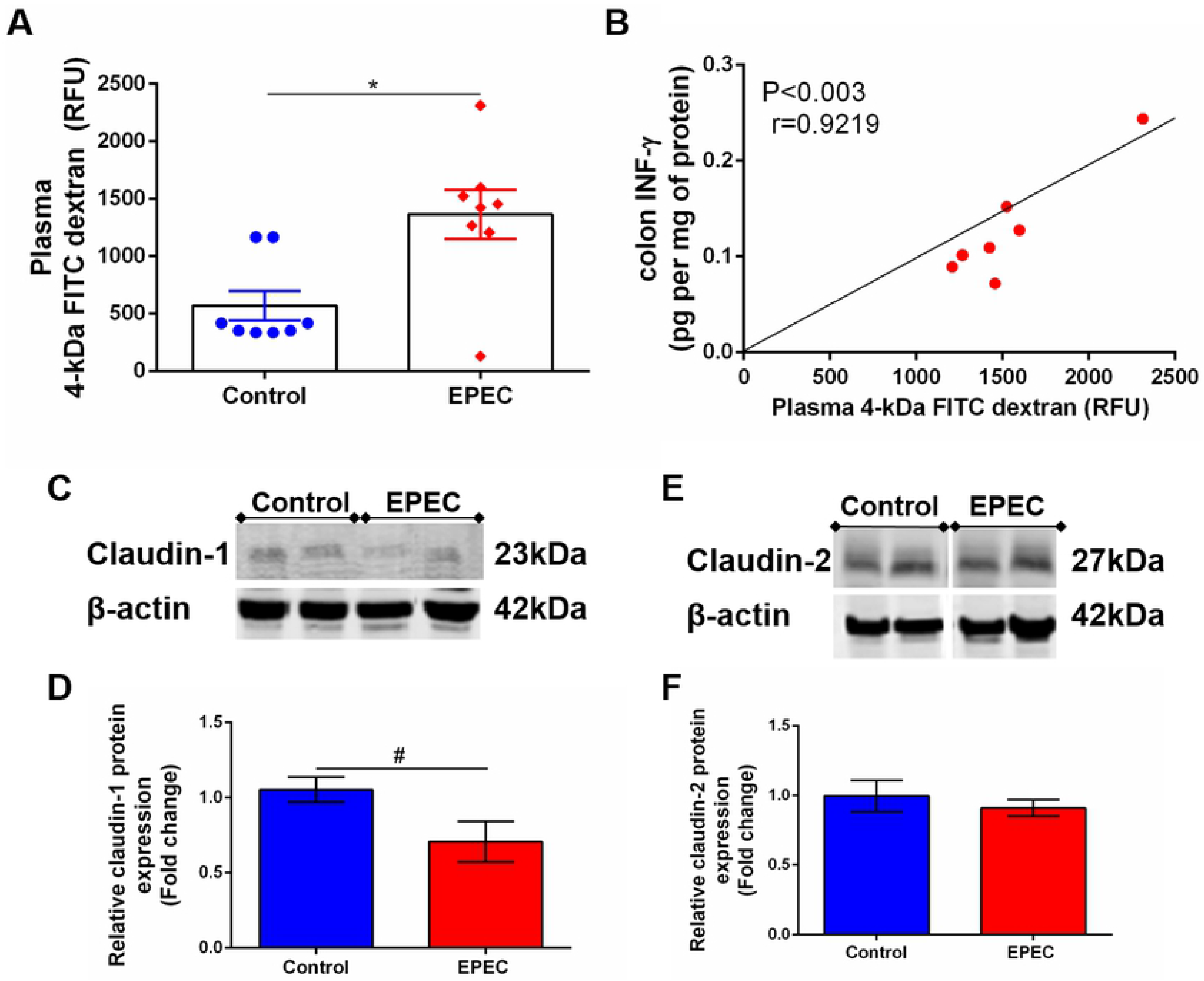
EPEC infection model increases intestinal permeability and decreases claudin-1 expression in colon of mice. (**A**) Intestinal permeability was determined in the plasma of control and EPEC-infected mice at day 7 p.i. using FITC dextran assay. Each point represents a mouse. Lines represent mean±SEM (n=8). *p<0.006 using Student’s *t-test.* (**B**) Significant positive correlation between diarrhea score and INF-γ levels in colon tissues of EPEC-infected mice at day 7 p.i. (Spearman rank test). (**C**) Representative western blots of levels of claudin-1 and β-actin in colonic tissues of control and EPEC-infected mice at day 7 p.i. (**D**) Quantification of western blot bands of claudin-1 and β-actin. Bars represent mean±SEM (n=3). #p<0.03 using Student’s *t-test.* (**E**) Representative western blots of levels of claudin-2 and β-actin in colonic tissues of control and EPEC-infected mice at day 7 p.i. (**F**) Quantification of western blot bands of Claudin-2 and β-actin. Bars represent mean±SEM (n=3).

Tight junctions play a crucial role in regulating intestinal permeability, we therefore, investigated if EPEC infection alters claudin-1 and claudin-2 expression in the colon of mice, and found that EPEC infection decreased claudin-1 (p<0.03, Fig 6C and 6D), but not claudin 2 (Fig 6E and 6F) in the colon when compared to control mice.

### Loss of escN expression in EPEC inhibits intestinal and systemic inflammation induced by wild-type EPEC infection model without changes in stools shedding in mice

The type 3 secretory system (T3SS) is essential for EPEC pathogenesis, and disruption of the *escN* gene (ATPase energizer), can lead to inefficient injection of EPEC into the host cell (Andrade et al., 2007). We therefore, tested whether inactivation of *escN* in EPEC strain could affect changes in body weight, intestinal and systemic inflammation in our EPEC infection model. The Δ*escN* EPEC-infected mice exhibited weight gain when compared to wild-type (WT) EPEC-infected mice on days 1 and 3 p.i (p<0.05 and p<0.001, respectively, Fig 7A). Deletion of Δ*escN* did not affect EPEC shedding in the stools (Fig 7B) and colonization intestinal tissues (Fig 7C). However, at day 8 p.i., tissue burden of Δ*escN* EPEC infected mice was detected only in the colonic tissue (p<0.01 Fig 7D). No histological changes were observed in the colon of mice infected with Δ*escN* EPEC when compared to mice infected with WT EPEC (Fig 7E). Furthermore, Δ*escN* EPEC infection resulted in decreased LCN-2 in the stools (day 2 and 7 p.i.) and cecal contents (day 3p.i.); and MPO in stools (day 2 p.i.), as well as IL-6 in ileum and colon (day 3 p.i. p<0.05, p<0.01 respectively, Fig 7H).

**Fig 7.**
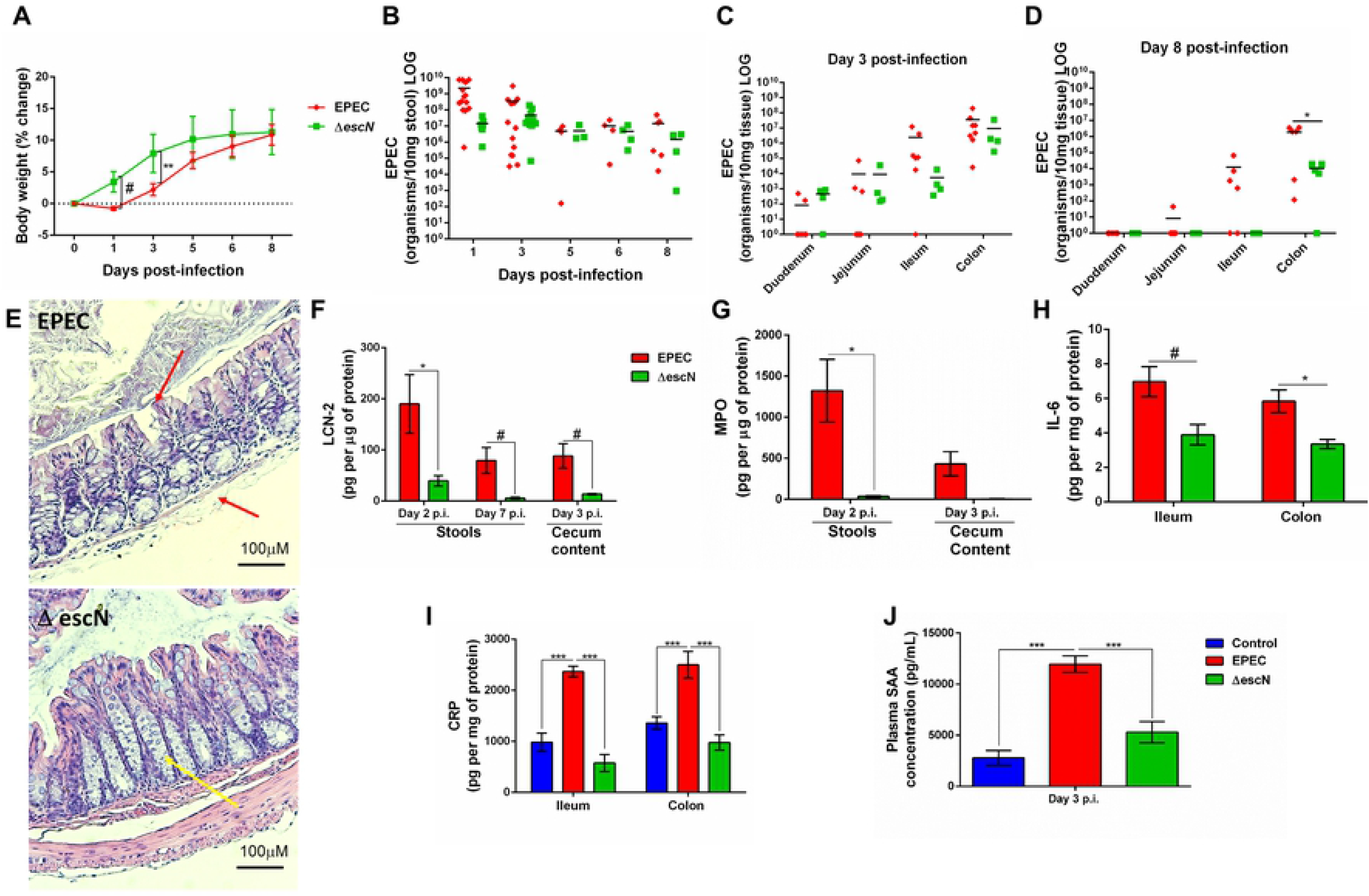
Inactivation of *escN* in EPEC decreases colonization in colon and reduces intestinal and systemic inflammation promoted by EPEC infection in mice. (**A**) Change in body weight of weaned C57BL/6 mice infected with 10^10^ CFU per mouse with WT EPEC (EPEC) or Δ*escN* EPEC *(ΔescN*, type 3 secretion system,T3SS, deficient) (n=12/group). Line graphs represents mean±SEM. **p<0.001 and #p<0.05 using multiple Student’s *t-test.* (**B**) Quantification of EPEC shedding in stools of WT EPEC (EPEC) or Δ*escN* EPEC infected mice. (**C-D**) Quantification of EPEC tissue burden in intestinal tissues (duodenum, jejunum, ileum and colon) at day 3 and 8 p.i. Bars represent mean±SEM. *p<0.01 using multiple Student’s *t-test.* (**E**) Representative H&E stains of colon from WT EPEC (EPEC) or Δ*escN* infected mice at day 3 p.i. Scale bars, 100μm. (**F**) LCN-2 levels measured in stools (day 2 and 7 p.i.) and cecal contents (day 3 p.i.) of WT EPEC (EPEC) or Δ*escN* infected mice. (**G**) MPO levels measured in stools (day 2 p.i.) and cecal contents (day 3 p.i.). (**H**) IL-6 levels measured in the ileal and colonic tissue lysates at day 3 p.i. using ELISA (**I**) CRP levels measured in the ileal and colonic tissue lysates of uninfected (control, in blue), WT EPEC (in red) or Δ*escN* EPEC (in green) infected mice at day 3 p.i. (**J**) Concentration of SAA in plasma at day 3 p.i. was determined using ELISA. Bars represent mean±SEM (n=8). # p<0.05, *p<0.01 and ***p<0.0001 using multiple Student’s *t-test* (F, G and H) or One-way ANOVA followed by Turkey’s test (I and J).

To assess whether WT EPEC infection induces systemic inflammation, as well as the participation of *escN* using Δ*escN* EPEC infection, CRP in intestinal tissues (ileum and colon) and SAA in plasma of mice collected at day 3 p.i. were measured. WT EPEC infection resulted in increased the levels of CRP in the intestinal tissues (p<0.0001, Fig 7I) and increased SAA (p<0.0001, Fig 7J) when compared to control mice. Whereas deletion of Δ*escN* in EPEC led to a significant decrease in levels of these markers of systemic inflammation at day 3 p.i. when compared to WT EPEC-infected mice (p<0.0001, Fig 6I and 6J).

Taken together, these findings showed that EPEC induces later colonic colonization, intestinal and systemic inflammation, in an *escN*-encoded T3SS-dependent manner, confirming the effectivity of this new EPEC infection model in mice.

## DISCUSSION

Typical EPEC infections have been suggested to be associated with inflammatory enteropathy and/or diarrhea in resource-limited populations [3, 44]. Infections caused by EPEC have been widely reported *in vitro* studies to demonstrate the effects of adherence traits [18] and type 3-secretion system (T3SS) [8]. However, there is still a need for a suitable *in vivo* model that clearly shows the effects of human EPEC infection in an intestinal environment. *Citrobacter rodentium* is a natural murine pathogen that has been used to study human EPEC and Enterohemorrhagic *E. coli* (EHEC) due to their genetic similarities; and has also been shown to cause A/E lesions, with the formation of pedestal structures and polarized actin accumulation at the site of infection [45, 46].

Here an EPEC murine infection model has been developed using mice pretreated antibiotic cocktail (vancomycin, gentamicin, metronidazole and colistin) to enable colonization, and induce growth impairment, acute diarrhea, intestinal damage and inflammation, as well as metabolomic perturbations and intestinal permeability alterations. Efforts of developing an EPEC model that mimics clinical outcomes, such as growth impairment and diarrhea as observed in humans has been a challenge for *in vivo* studies [20, 25, 30, 43, 47]. Most of the previous EPEC infection murine models have used streptomycin in order to disrupt the intestinal microbiota and promote colonization in mice [27, 28, 30, 31, 46]. Recently, a study demonstrated that mice are susceptible to EPEC colonization in an age and microbiota disruption-dependent manner, with the infant mice being more susceptible [30]. We have previously shown that disruption of intestinal microbiota using a broad-spectrum antibiotic cocktail enabled colonization of bacterial pathogens such as ETEC [48], *C. jejuni* [33] and *S. flexneri* [34] in C57BL/6 mice with mice developing diarrhea. This current study also used the same antibiotic cocktail to enable the assessment of disease outcomes as a result of EPEC infection.

We also identified a two-phase disease promoted by the EPEC infection model, an acute symptomatic and a later asymptomatic phase. In the acute symptomatic phase of the disease, EPEC-infected mice had growth impairment and diarrhea as clinical outcomes, and this was accompanied with ileal and colonic damage (loss of integrity of epithelial cells, edema in submucosa and intense infiltrate of inflammatory cells) and intense inflammation. Similar findings showing disruption of colonic damage following EPEC infection has also been previously reported [31].

In humans, EPEC infection promotes watery diarrhea and dehydration [5]. Most of the EPEC infected mice developed moderate to severe diarrhea at day 3 p.i. Previously, C57BL/6 mice infected with EPEC have been reported to develop semi-solid stools in the proximal colon with no apparent diarrhea [25].

MPO has been used as biomarker of enteropathy in clinical studies [36, 49], also exhibiting inflammation, growth and development decrements in children infected with different enteropathogens in low-income countries. In our present study, MPO was increased during acute phase of disease, in the intestinal tissues (ileum and colon); MPO was detected mainly in stools of mice that developed soft or unformed stools or diarrhea. In addition, LCN-2 which is known as a neutrophil gelatinase-associated protein expressed by intestinal epithelial cells [50], was detected in higher concentrations in stools of all the mice that were infected with EPEC. Particularly impressive in our model was the striking inflammatory enteropathy (as evidenced by fecal LCN-2 and MPO during acute infection) with EPEC infected mice developing watery mucoid stools.

EPEC infection in infant C57BL/6 mice has been previously reported to colonize the small intestine and colon for 3 days in a study of as a result of human milk oligosaccharides administration [29]. Here, the ileum and colon were markedly colonized by EPEC during infection challenge. Similar to our findings, EPEC infection in other mouse models have also been reported to colonize the ileal and colonic tissues [25, 30]. Moreover, the colon in our model was most affected by EPEC infection with higher colonization by EPEC; and similar findings have been previously reported in germ-free mice infected with EPEC [30].

The pro-inflammatory cytokines such as IL-6, IL-1β, IL-23, IL-22 and INF-γ were increased in colon of mice infected with EPEC. IL-6 is a pleiotropic cytokine showing a pro-inflammatory phenotype and is protective against infection. For instance, deficiency of IL-6 in C57BL/6 mice has been reported to cause colonic damage, increase infiltrate of inflammatory cells and apoptosis during infection with *C. rodentium* [51]. Mice lacking IL-1β are more susceptible to *C. rodentium-induced* colonic inflammation [52], in contrast blocking IL-1β in EPEC-infected mice with persistent IL-1β response decreased the colonic damage [53], suggesting the role of IL-1β during intestinal infection in a concentration- and timing-dependent manner. Binding of IL-1β, to IL-1 receptor type I (IL-1RI) and activation of nuclear factor κB (NFκB), promotes the recruitment of inflammatory cells at the site of inflammation by inducing the expression of adhesion molecules on endothelial cells and the release of chemokines [54, 55]. In our model, EPEC infection increased the expression of adhesion molecules (VCAM1, ICAM1 and SELP), as well as chemokines (CCL2, CCL5, CCL19, CXCL10 and CXCL11) and the chemokine receptors CCR2 (activated by CCL2 and expressed by macrophage and lymphocytes) and CCR7 (activated by CCL19, promoting migration of dendritic cells, monocytes and T cells) [56], contributing to the intense recruitment of inflammatory cells in the colon, but not in the ileum, of EPEC-infected mice.

Similar to IL-6, INF-γ is a pleiotropic protein that promotes the transcription of pro-inflammatory mediators and CXCL10 (by binding to CXCR3 in order to promote the recruitment of monocytes/macrophages and T cells at the site of infection), [57] and were both upregulated in the colon of mice following challenge with EPEC, likely activating STAT1 by binding to INF-γ receptor (INFGR) [58]. In fecal samples from children with symptomatically EPEC infection, intermediate levels of INF-γ have been associated with an increase in infection duration [59]. INF-γ levels were increased in colonic tissues in our study, and severity of diarrhea was associated with higher levels of INF-γ in EPEC-infected mice at day 3 p.i.

IL-23, was also increased following EPEC infection, and has been shown to be required to promote IL-22 expression, a cytokine involved in promoting tissue regeneration and regulating inflammation, and also to negatively control the potentially deleterious production of IL-12 [60]. The data therefore, suggests that an increase in these cytokines during EPEC-infection in our model is protective, but not enough to prevent the intestinal damage promoted by EPEC. In *C. rodentium* infection, lacking of IL-23 in macrophages led to increased mortality in mice [60]. Here, we also provided data suggesting that our EPEC infection model was able to activate NFκB via IL-1β, STAT1 via INF-γ, and STAT3 via IL-22 and IL-6, but not CREB. STAT-1 and STAT-3 contribute to the expression of pro-apoptotic and anti-apoptotic genes respectively [57, 61]. However, in the present study, it seemed that the response mediated by STAT-1 (whose expression was higher than STAT3 in colon of EPEC-infected mice) prevailed over anti-apoptotic response promoted by STAT3, once increased cleaved caspase, a marker of apoptosis, were detected. Moreover, because our EPEC infection model exhibits evident diarrhea more investigation of how these cytokines contribute to its pathogeneses is needed.

Furthermore, during acute phase of infection, EPEC infection resulted in perturbations of multiple biochemical pathways, with the TCA cycle intermediates appearing to be the most sensitive to EPEC infection. The TCA cycle in *E. coli* is linked to energy metabolism in which CO_2_ concomitant is oxidized from pyruvate leading to production of NADH and FADH_2_ [62]. In our model, the TCA cycle metabolites were excreted in lower quantities following EPEC infection, suggesting that energy production was reduced or conserved in the infected host. A shut-down of the TCA cycle during infection suggests that the energy requirements of the host were not met, potentially explaining in part the significant weight loss in the infected mice. *C. jejuni* infection in zinc deficient mice have also been reported to perturb the TCA cycle, affecting amino acid and muscle catabolism as a result of increased creatine excretion [33]. Pantothenate is the key precursor of the fundamental TCA cycle cofactor, coenzyme A [63]. Reduced pantothenate excretion following EPEC colonization further adds to the TCA cycle disruption by infection. Interestingly, excretion of creatine which is a source for energy production in the form of ATP was also increased during infection. Sugiharto and colleagues reported on post-weaning pigs infected with *E. coli* F18 and found that there was a reduction in creatine and betaine which was due to inhibition of antioxidant system that resulted in piglets developing diarrhea [64].

Taurine has been shown to possess antioxidant properties and its concentrations are elevated in inflamed tissues where oxidants are abundant [65, 66]. EPEC infection in this study was characterized by elevated urinary taurine excretion. As we have previously observed [67], treating rodents with antibiotics suppresses the bacterial metabolism of taurine thus increasing taurine bioavailability and uptake in the host reflected by greater urinary taurine excretion.

Metabolites derived from bacteria in the gut were excreted in greater amounts following infection suggesting gut microbial metabolism was altered by EPEC infection. These findings help to understand host metabolism during infection, suggesting potential pathways to be further explored and targeted in future studies.

In relation to later phase of EPEC infection, we observed an increase in TNF-α in the colon and ileum, as well as increased LCN-2 in stools samples and increased intestinal permeability and decreased claudin-1. Despite TNF-α gene expression, as well as its receptor TNFRS being increased during the acute phase of disease, the TNF-α protein levels were increased in intestinal tissues of EPEC-infected mice only during the later phase. TNF-α synthesis is promoted by NFκB activation which in turn can be promoted by IL-1β [68]. The biological effects of TNF-α mediated by binding to TNFRS include inflammation, apoptosis and tissue regeneration via activation of NFκB, caspase-8 and AKT respectively [68]. Similar to our findings, later increases of TNF-α in the ileum and colon has been observed by others in EPEC-infected mice at day 5p.i. [69], findings that we showed at day 7 p.i. This increase in TNF-α was associated with increased LCN-2 in stools, indicating the presence of intestinal damage and inflammation, despite partial recovery from the acute phase of disease. TNF-α has been shown to induce LCN-2 expression by activating NFκB [50]. TNF-α and INF-γ have been associated with a loss of integrity of the intestinal epithelial barrier [70]. Despite TNF-α, but not INF-γ, being increased in colonic tissues in EPEC-infected mice, only INF-γ levels were associated positively with an increase in intestinal permeability; a similar association has also been reported in an *in vitro* study using T84 epithelial cell monolayer [71]. The permeability of the intestinal barrier is regulated by the tight junction proteins [72]. EPEC infection has been reported to impair tight junction barrier function of ileal and colonic mucosa [27, 31, 46, 73, 74]. Claudin-1, a component of tight junction expressed by epithelial cells from small and large intestine, is responsible for increasing barrier tightness [75]. In our model, a decrease of claudin-1 in the colon of EPEC-infected mice was associated with an increase in intestinal permeability. Although we detected alterations in intestinal permeability at the later stage of EPEC infection, this might have been due in part to an increase in systemic markers (SAA and CRP) that were detected at day 3 p.i. during the acute phase leading to disruption of intestinal tight junctions. SAA has been a biomarker of enteropathy in clinical studies also associated with inflammation, and with growth and developmental impairment in children infected with multiple enteropathogens in low-income countries [49].

The T3SS is essential for EPEC pathogenesis and requires an effective ATPase energizer, *escN* [8, 76, 77]. Here, we also demonstrated that mice infected with *escN* deletion mutant resulted in diminished growth impairment and inflammation. Even without an effective T3SS, *escN* mutant was able to colonize all sections of the intestinal tissue, albeit at much lower levels, to day 8 pi, as shown by our results. These findings reinforce the importance of a functional T3SS in the virulence of EPEC in this model. [6, 8]

In conclusion, our findings showed that EPEC infection causes growth impairment, diarrhea and increased inflammatory responses in weaned antibiotic pretreated mice. These effects were also dependent on an intact EPEC T3SS. In addition, metabolic perturbations and intestinal permeability were also observed in mice with EPEC infection, suggesting relevant biochemical pathways involved. Further, the findings presented here suggest that EPEC infections leads to an increase in intestinal and systemic inflammatory responses and transient overt diarrhea and growth impairment, as is often seen in children with EPEC infections. This EPEC infection model also presents two phases of diseases: an acute symptomatic and a later asymptomatic phase (Fig 8). This model can help further explore mechanisms involved in EPEC pathogenesis and perhaps facilitate the development of vaccines or therapeutic interventions.

**Fig 8.**
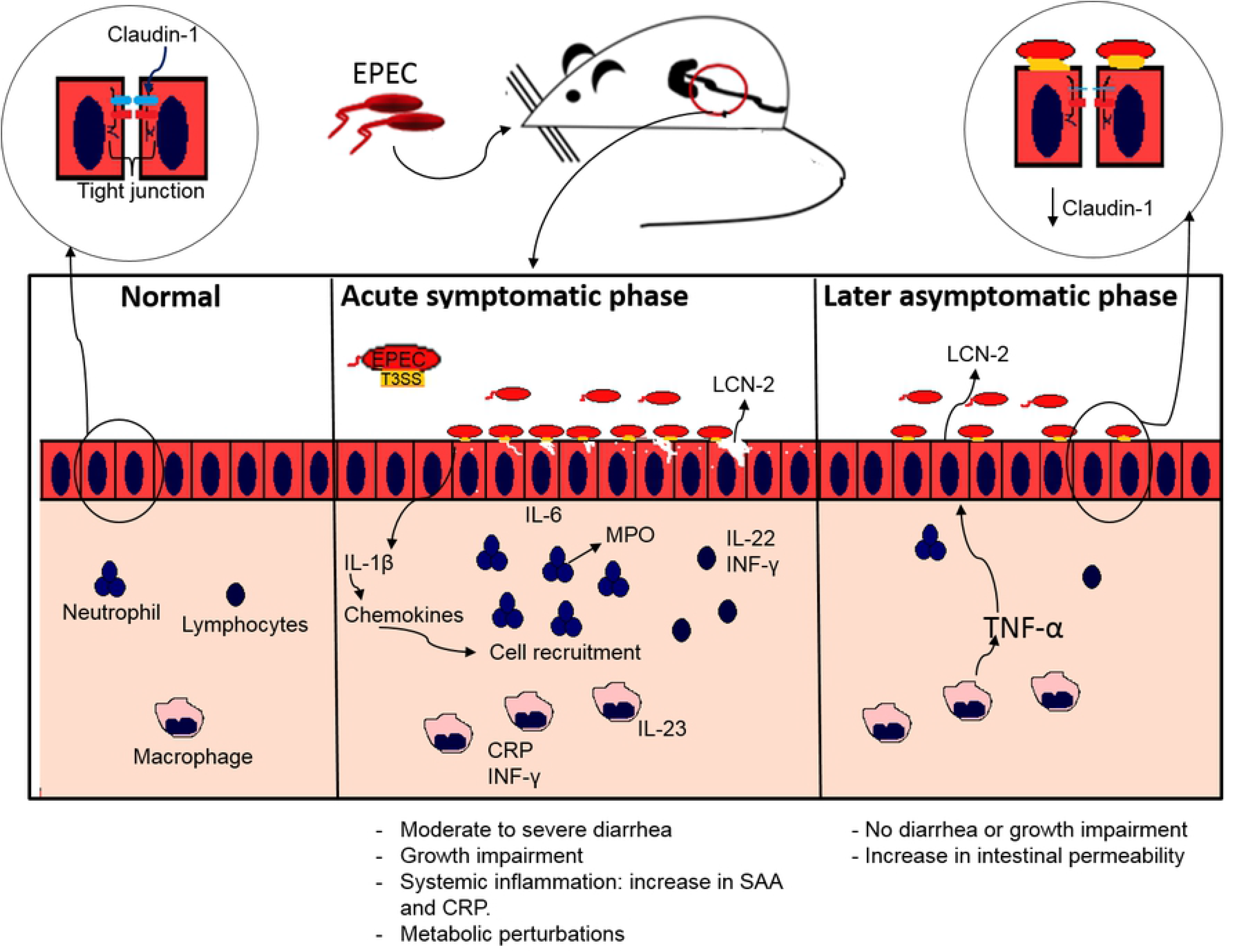
Proposed model of EPEC infection antibiotic pretreated mice displaying acute symptomatic and late asymptomatic phase. During acute symptomatic phase, colonization by EPEC leads to growth impairment, accompanied with moderate to severe diarrhea. Adherence of EPEC on the intestinal epithelial cells leads to increased IL-1β, which in turn stimulates chemokines synthesis, and consequently recruitment of neutrophils, macrophages and lymphocytes, resulting in increased release of MPO by neutrophils and LCN-2 by epithelial cells, as well as pro-inflammatory (IL-6, IL-23, INF-γ) and IL-22 cytokines indicating intestinal inflammation. A further increase in CRP and SAA indicates acute systemic inflammation. During the later asymptomatic phase an increase in LCN-2 inflammatory biomarker and an increase in TNF-α leads to increased intestinal permeability affecting the tight junction integrity by decreasing claudin-1 with no signs of diarrhea and growth impairment.

## MATERIALS AND METHODS

### Ethics statement

The mice used in the study have been handled with strict accordance with the recommendations in the Guide for the Care and Use of Laboratory Animals of the National Institutes of Health. The protocol has been approved by the Committee on the Ethics of Animal Experiments of the University of Virginia (Protocol Number: 3315). All efforts were made to minimize suffering. This is also in accordance with the Institutional Animal Care and Use Committee policies of the University of Virginia. The University of Virginia is accredited by the Association for the Assessment and Accreditation of Laboratory Animal Care, International (AAALAC).

### Mice

Mice used in this study were male, 22 days old, C57BL/6 strain, ordered from Jackson Laboratories (Bar Harbor, ME). Mice weighed approximately 15 grams on arrival and were co-housed in groups of up to 4 animals per cage. The vivarium was kept at a temperature of between 20-23 °C with a 14-hour light and 10-hour dark cycle. Mice were allowed to acclimate for 3 days upon arrival. Mice were fed standard rodent House Chow diet (HC) from arrival and throughout the infection challenge.

### EPEC inoculum preparation

Bacterial strains used included: wild type EPEC E2348/69 [78] and EPEC E2348/69 Δ*escN* CVD425 [79]. Bacterial cultures were prepared from glycerol stocks maintained at −80 °C. Cultures were grown in 20 mL Dulbecco’s modified Eagle’s medium containing phenol red (DMEM) at 37 °C in a shaking incubator until cultures turned orange indicating optimal growth, OD_600_ ~ 0.6. Cultures were centrifuged at 3500 x g for 10 min at 4 °C. The bacterial pellet was resuspended in DMEM high glucose in order to obtain 10^10^ CFU/mL.

### EPEC infection model

Four days prior to challenge with EPEC, mice were given an antibiotic cocktail of gentamicin (35 mg/L), vancomycin (45 mg/L), metronidazole (215 mg/L), and colistin (850 U/ml) in drinking water for 3 days in order to disrupt resident microbiota followed by 1 day on normal water in order to clear antibiotics [48]. Then, mice were administered 100 μL of 10^10^ CFU/mL (10^10^ bacteria per mouse) bacterial culture in DMEM high glucose orally using 22-gauge feeding needles. Uninfected control mice were administered only 100μl DMEM high glucose.

After infection, all mice were weighed and stools were collected daily until 8-days post infection (p.i.). Mice were euthanized on days 3, 7 and 8 p.i. Fig. 1A shows the schematic presentation summarizing the experimental procedure.

### Analysis of clinical outcomes

The clinical outcomes, body weight and diarrhea, were assessed daily. Body weight was measured for 9 days (starting before infection, day 0) and percentage of changes in body weight was measured based on each individual mouse weight from day 0 before infection. Diarrhea scores were measured until day 7 p.i. Diarrhea score were based on the following 0 to 4: 0-well-formed pellets; 1-stick stools adhering in microtubes wall; 2-pasty stools with or without mucus; 3-watery stools with or without mucus; and 4-Stools with blood.

### Tissue burden and Stool shedding

For stool shedding, DNA was extracted from stools using the QIAamp DNA Stool Mini kit (Qiagen) according to the manufacturer’s instructions. For tissue burden, tissues were homogenized using the beat-beater and DNA was extracted using DNeasy Kit (Qiagen) according to the manufacturer’s instructions. The *eae* (intimin) gene was used as a specific target for detecting EPEC in stools and tissues. Primer sequences included *eae* 5’-CCCGAATTCGGCACAAGCATAAGC-3’ (sense) and 5’-CCCGGATCCGTCTCGCCAGTATTCG-3’ (antisense) [80]. Real-time PCR was performed using Bio-Rad CFX under the following conditions: 95 °C for 3 min, followed by 40 cycles of 15 sec at 95 °C 60 sec at 55 °C and lastly 20 sec at 72 °C.

### EPEC adherence on the intestine

Ileal tissue segments from EPEC-infected mice on day 3 p.i. were fixed in 4% formalin, embedded in paraffin, the slides were stained with anti-rabbit intimin at the University of Virginia Histology core, and viewed using light microscope.

### Transmission Electron Microscopy (TEM)

For TEM, the ileal tissues from EPEC-infected mice on day 3 p.i. and uninfected (control) were fixed with 4% glutaraldehyde.

The samples were washed with 1X cacodylate buffer for 10 min and placed in 2% osmium tetroxide for 1 hour. Then washed for 10 min with cacodylate buffer and distilled water. Followed by dehydration with 30% ethanol for 10 min and concentrations of 50%, 70%, 95% and 100% ethanol all for 10 min each. About 1:1 ethanol/propylene oxide was used for 10 min followed by 100% propylene oxide (PO) for 10 min. The samples were then placed in 1:1 of PO/epoxy resin (EPON) overnight followed by 1:2 PO/EPON for 2 hours, then 1:4 PO/EPON for 4 hours and lastly 100% EPON for overnight. The samples were then embedded in fresh 100% EPON and allowed to bake in a 65 °C oven. Ultra-thin sections were cut at 75 nm and picked up on 200 mesh copper grids. Sections were stained with 0.25% lead citrate and 2% uranyl acetate. The slides were viewed using JEOL 1230 microscope, with 4k x 4k CCD camera.

### Histology analysis

Mice intestinal samples (ileum and colon) from day 3 or day 7 p.i. were fixed in 10% neutral buffered formalin for 20h, dehydrated and embedded in paraffin. Ileal and colonic sections (5 μm) were stained with hematoxylin and eosin staining (H&E) and were examined using light microscopy. Histological damage scores were determined by quantifying the intensity of epithelial tissue damage (0-3, 0-no damage, 1-mild, 2-moderate, 3-extensive), edema in submucosa layer (0-3) and neutrophil infiltration (0-3).

### Intestinal inflammatory response by Enzyme-linked immunosorbent assay (ELISA)

Protein lysates were extracted from the stools, ileum, colon and cecal contents using radioimmunoprecipitation assay (RIPA) buffer (20 mM Tris [pH 7.5], 150 mM NaCl, 1% Nonidet P-40, 0.5% sodium deoxycholate, 1 mM EDTA, 0.1% SDS) containing protease inhibitor cocktail (Roche) and phosphatase inhibitors (1 mM sodium orthovanadate, 5 mM sodium fluoride, 1 mM microcystin LR, and 5 mM ß-glycerophosphate). Lysates were centrifuged at 13000 rpm for 15 min and the supernatant was used to perform the protein assay using the bicinchoninic acid assay (Thermo Fisher Scientific). Inflammatory biomarkers (LCN-2, MPO, IL-23, IL-22, IL-17, GMCSF, IL-33 and IL-10) were measured using a commercial ELISA kit (R&D Systems) according to the manufacturer instructions. Interleukin-6 (IL-6), IL-1β, interferon gamma (INF-γ), TNF-α, KC (analogue to human IL-8), IL-18, and CRP levels were measured using a ProcartPlex multiplex immunoassay (Invitrogen) by Luminex (Biorad). Biomarkers levels were measured as picograms per milligram of protein.

### Phosphorylated-STAT3, phosphorylated-CREB and cleaved caspase-3 measurement

Colonic samples from EPEC-infected and control mice collected at day 3 p.i. were homogenized in ice-cold using RIPA buffer containing protease and phosphatase inhibitors. The colonic levels of phosphorylated STAT3 3 (pSTAT3), phosphorylated CREB (pCREB) and cleaved caspase-3 were measured using ELISA kits (R&D Systems) according to the manufacturer’s instructions.

### TaqMan-real time polymerase chain reaction (qPCR)

The isolation of total RNA from colon tissues of EPEC-infected (presenting moderate or severe diarrhea) and control mice were performed by using a Qiagen RNeasy mini kit and QIAcube. cDNA was synthetized from 1μg of total RNA, quantified by Qubit 3 fluorometer 3000 (Invitrogen) and purified by deoxyribonuclease I (Invitrogen) treatment, with the iScript cDNA (Bio-Rad) as described by manufacturer instructions. qPCR was performed with 50 ng of cDNA in each well and SensiFAST probe no-ROX mix (Bioline) using a CFX Connect system (Bio-Rad) with the following conditions: 95 °C for 2 min, 40 cycles of 95 °C for 10 s and 60°C for 50 s. A pre-designed TaqMan array mouse immune fast 96-well plates (Applied Biosystems) was used to assess the expression of 92 genes listed in supplementary Table 1. Glyceraldehyde-3-phosphate dehydrogenase (GAPDH) was used as a reference gene. All fold changes were determined using the ΔΔC_t_ method [81].

### *In vivo* intestinal permeability assay

For assessing *in vivo* intestinal permeability fluorescein isothiocyanate (FITC)-labeled dextran assay (4kDa, Sigma Aldrich) was used. Mice were deprived food, with free access to water, for 4 h. Then, 200 μL of FITC-dextran solution (80mg/mL in water) was administered by oral gavage for each mouse. After 4 h of FITC administration, mice were anesthetized to collect blood using cardiac puncture. Then, the blood samples were centrifuged (5 min, 8000 rpm, 4 °C) and plasma was obtained. Fluorescence intensity in 100 μL of plasma placed on Qubit 0.5 mL-microtubes (Life Technologies) was measured using Qubit 3fluorometer (Life Technologies) using an excitation wavelength of 470 nm. A plasma sample from mice not receiving FITC-dextran solution was used as a blank.

### ^1^H NMR spectroscopy based metabolic profiling

Urine specimen were collected in a sterile 1.5 mL eppendorf tube and placed at −80 °C until further analysis. The metabolic profiling was performed on all urine samples using ‘H nuclear magnetic resonance (NMR) spectroscopy. A 30 μL urine aliquot was combined with 30 μL of phosphate buffer (pH 7.4, 100% D_2_O, 0.2 M Na_2_HPO_4_/NaH_2_PO_4_) containing 1 mM of the internal standard, 3-(trimethylsilyl)-[2,2,3,3^-2^H_4_]-propionic acid (TSP) and 2 mM sodium azide (NaN_3_) as a bacteriocide. Samples were vortexed and spun at 13,000 x g for 10 min and 50 μL of the supernatant was then transferred to 1.7 mm NMR tubes. Spectroscopic analysis was performed on a 600 MHz Bruker Avance^TM^ NMR spectrometer at 300 K using a Bruker BBI probe and an automated SampleJet for tube handling (Bruker, Germany). ? NMR spectra of the urine samples were acquired using a standard one-dimensional pulse sequence [recycle delay (RD) −90°-t_1_-90°-t_m_-90°-acquire free induction decay (FID)]. The water signal was suppressed through irradiation during the RD of 4 s and a mixing time of (t_m_) 100 ms was used. For each spectrum, 64 scans were obtained into 64 K data points using a spectral width of 12.001 ppm. The NMR spectra were calibrated to the TSP resonance at 0 ppm using TopSpin 3.5 NMR software (Bruker, Germany) and imported into MATLAB (R2018a, Mathworks Inc, Natwick, MA) using in-house scripts. Regions containing the TSP, water and urea resonances were removed from the urinary spectra. ^1^H NMR spectra were manually aligned and normalized to the unit area.

### Western blotting

Colon tissues from EPEC-infected and control mice at day 7 p.i. were collected, lysed using RIPA lysis buffer containing complete EDTA-free protease inhibitor cocktail (Roche) and phosStop (Roche) and centrifuged (17 min, 4°C, 13000 rpm). Then the supernatant was collected for extracting protein. Protein concentrations were determined through the bicinchoninic acid assay according to the manufacturer’s protocol (Thermo Fisher Scientific). Reduced 60 μg protein samples (previously prepared with sample reducing agent-Invitrogen- and protein loading buffer-LI-COR) were denatured at 95 °C for 5 min, separated on NuPAGE 4-12 % BIS-Tris gel (Invitrogen) and transferred to nitrocellulose membranes (Life Technologies) for 2 h. The membranes were then immersed in iBind fluorescent detection solution (Life technologies) and placed in a iBIND automated Western device (Life Technologies) overnight at 4 °C for blocking, incubating with primary antibodies (rabbit anti-β-actin, 1:1000, Thermo Fisher Scientific, PA1-183; mouse anti-claudin-2, 3:500, Invitrogen, 325600; rabbit anti-claudin-1, 1:1000, Novus biological, NBP1-77036) and secondary antibodies (Cy3-conjugated AffiniPure donkey anti-rabbit, 711-165-152, 1:1000, Jackson ImmunoResearch and Cy5-conjugated AffiniPure donkey anti-mouse, 715-175-150, 1:1000, Jackson ImmunoResearch). Then, the membranes were immersed in ultrapure water and fluorescent signal was detected using the Typhoon system (GE healthcare). Densitometric quantification of bands was performed using ImageJ software (NIH, Bethesda, MD, USA).

### Systemic inflammation analysis

Blood collected at day 3 p.i. was centrifuged at 8000 rpm for 5 min at 4 °C in order to obtain the plasma, to measure the levels of SAA as a marker of systemic inflammation. The levels of SAA were measured using a commercial ELISA kit (R&D Systems) according to the manufacturer’s instructions. The results were expressed as picograms per milliliter.

### Statistical analysis

All data were analyzed using GraphPad Prism 7 software (GraphPad Software). Data are presented as the mean ± standard error of the mean (SEM) or as medians when appropriate. Student’s t test and one-way Analysis of Variance (ANOVA) followed by Tukey’s *test* were used to compare means, and the Kruskal-Wallis and Dunn tests were used to compare medians. Spearman rank test was used to correlation analyses. Differences were considered significant when p < 0.05. Experiments were repeated at least two times.

For metabolomics data analysis multivariate statistical modelling was performed including principal component analysis (PCA) using the Imperial Metabolic Profiling and Chemometrics Toolbox (https://github.com/csmsoftware/IMPaCTS) in MATLAB (Version 2018a, Mathworks Inc) and unsupervised hierarchical clustering analysis (HCA) to unveil metabolic differences between EPEC infected and control mice. Unsupervised clustering for all samples was done with the use of the normalized abundance of metabolites that were identified through the PCA models. Hierarchical clusters were generated with the use of an average-linkage method by means of the pdist and linkage functions in the MATLAB bioinformatics toolbox. Heat maps and dendrograms after the HCA were generated with MATLAB imagesc and dendrogram functions, respectively. Pathway analysis was performed using the MetaboAnalyst 4.0 platform (https://www.metaboanalyst.ca/).

## ACKNOWLEDGEMENTS

Research reported in this publication was supported by the Bill & Melinda Gates Foundation Opportunity ID: OPP1137923, the NIH’s Fogarty International Center award number D43TW006578, NIAID award number U19AI109776 and National Research Foundation (NRF). The content is solely the responsibility of the authors and does not necessarily represent the official views of the funders.

## AUTHOR CONTRIBUTION

SEL, DTB, RLG, JPN, NP and ANTH designed the project. SEL, DVS and DTB performed the experiments. SEL, DVS, DTB, and RLG analyzed the data and wrote the manuscript. PHQSM assisted in analysis of results. NG and JRS performed and assisted with metabolome analysis. All authors revised the manuscript. NP,ANTH, RLG, JPN supervised the project.

## CONFLICTS OF INTEREST

The authors declare no financial and non-financial competing interest.

## REFERENCES

1. Liu L, Oza S, Hogan D, Perin J, Rudan I, Lawn JE, et al. Global, regional, and national causes of child mortality in 2000–13, with projections to inform post-2015 priorities: an updated systematic analysis. Lancet. 2015;385(9966):430–40. doi: 10.1016/S0140-6736(14)61698-6. PubMed PMID: 25280870.

2. Kotloff KL, Nataro JP, Blackwelder WC, Nasrin D, Farag TH, Panchalingam S, et al. Burden and aetiology of diarrhoeal disease in infants and young children in developing countries (the Global Enteric Multicenter Study, GEMS): a prospective, case-control study. Lancet. 2013;382(9888):209–22. Epub 2013/05/18. doi: 10.1016/S0140-6736(13)60844-2. PubMed PMID: 23680352.

3. Platts-Mills JA, Babji S, Bodhidatta L, Gratz J, Haque R, Havt A, et al. Pathogen-specific burdens of community diarrhoea in developing countries: a multisite birth cohort study (MAL-ED). The Lancet Global health. 2015;3(9):e564–75. Epub 2015/07/24. doi: 10.1016/S2214-109X(15)00151-5. PubMed PMID: 26202075.

4. Kaper JB, Nataro JP, Mobley HL. Pathogenic Escherichia coli. Nat Rev Microbiol. 2004;2(2):123–40. Epub 2004/03/26. doi: 10.1038/nrmicro818. PubMed PMID: 15040260.

5. Guerrant RL, Walker DH, Weller PF. Tropical infectious diseases: principles, pathogens and practice. Third edition. ed. Edinburgh: Saunders/Elsevier; 2011. xxiv, 1130 pages p.

6. Frankel G, Phillips AD, Rosenshine I, Dougan G, Kaper JB, Knutton S. Enteropathogenic and enterohaemorrhagic Escherichia coli: more subversive elements. Mol Microbiol. 1998;30(5):911–21. PubMed PMID: 9988469.

7. Friedberg D, Umanski T, Fang Y, Rosenshine I. Hierarchy in the expression of the locus of enterocyte effacement genes of enteropathogenic Escherichia coli. Mol Microbiol. 1999;34(5):941–52. Epub 1999/12/14. PubMed PMID: 10594820.

8. Andrade A, Pardo JP, Espinosa N, Perez-Hernandez G, Gonzalez-Pedrajo B. Enzymatic characterization of the enteropathogenic Escherichia coli type III secretion ATPase EscN. Arch Biochem Biophys. 2007;468(1):121–7. doi: 10.1016/j.abb.2007.09.020. PubMed PMID: 17964526.

9. Giron JA, Ho AS, Schoolnik GK. An inducible bundle-forming pilus of enteropathogenic Escherichia coli. Science. 1991;254(5032):710–3. Epub 1991/11/01. PubMed PMID: 1683004.

10. Knutton S, Baldwin T, Williams PH, McNeish AS. Actin accumulation at sites of bacterial adhesion to tissue culture cells: basis of a new diagnostic test for enteropathogenic and enterohemorrhagic Escherichia coli. Infect Immun. 1989;57(4):1290–8. Epub 1989/04/01. PubMed PMID: 2647635; PubMed Central PMCID: PMC313264.

11. Scaletsky IC, Aranda KR, Souza TB, Silva NP. Adherence factors in atypical enteropathogenic Escherichia coli strains expressing the localized adherence-like pattern in HEp-2 cells. Journal of clinical microbiology. 2010;48(1):302–6. Epub 2009/10/30. doi: 10.1128/JCM.01980-09. PubMed PMID: 19864474; PubMed Central PMCID: PMCPMC2812252.

12. Pelayo JS, Scaletsky ICA, Pedroso MZ, Sperandio V, Giron JA, Frankel G, et al. Virulence properties of atypical EPEC strains. J Med Microbiol. 1999;48(1):41–9. Epub 1999/01/27. doi: 10.1099/00222615-48-1-41. PubMed PMID: 9920124.

13. Mora A, Blanco M, Yamamoto D, Dahbi G, Blanco JE, Lopez C, et al. HeLa-cell adherence patterns and actin aggregation of enteropathogenic Escherichia coli (EPEC) and Shiga-toxin-producing E. coli (STEC) strains carrying different eae and tir alleles. Int Microbiol. 2009;12(4):243–51. Epub 2010/01/30. PubMed PMID: 20112229.

14. Abe CM, Trabulsi LR, Blanco J, Blanco M, Dahbi G, Blanco JE, et al. Virulence features of atypical enteropathogenic Escherichia coli identified by the eae(+) EAF-negative stx(-) genetic profile. Diagn Microbiol Infect Dis. 2009;64(4):357–65. Epub 2009/05/16. doi: 10.1016/j.diagmicrobio.2009.03.025. PubMed PMID: 19442475.

15. Hu J, Torres AG. Enteropathogenic Escherichia coli: foe or innocent bystander? Clin Microbiol Infect. 2015;21(8):729–34. Epub 2015/03/03. doi: 10.1016/j.cmi.2015.01.015. PubMed PMID: 25726041; PubMed Central PMCID: PMCPMC4497942.

16. Freter R, Brickner H, Fekete J, Vickerman MM, Carey KE. Survival and implantation of Escherichia coli in the intestinal tract. Infect Immun. 1983;39(2):686–703. Epub 1983/02/01. PubMed PMID: 6339389; PubMed Central PMCID: PMCPMC348005.

17. Fabich AJ, Jones SA, Chowdhury FZ, Cernosek A, Anderson A, Smalley D, et al. Comparison of carbon nutrition for pathogenic and commensal Escherichia coli strains in the mouse intestine. Infect Immun. 2008;76(3):1143–52. Epub 2008/01/09. doi: 10.1128/IAI.01386-07. PubMed PMID: 18180286; PubMed Central PMCID: PMCPMC2258830.

18. Knutton S, Shaw RK, Anantha RP, Donnenberg MS, Zorgani AA. The type IV bundle-forming pilus of enteropathogenic Escherichia coli undergoes dramatic alterations in structure associated with bacterial adherence, aggregation and dispersal. Mol Microbiol. 1999;33(3):499–509. PubMed PMID: 10417641.

19. Leverton LQ, Kaper JB. Temporal expression of enteropathogenic Escherichia coli virulence genes in an in vitro model of infection. Infect Immun. 2005;73(2):1034–43. Epub 2005/01/25. doi: 10.1128/IAI.73.2.1034-1043.2005. PubMed PMID: 15664947; PubMed Central PMCID: PMC546935.

20. Law RJ, Gur-Arie L, Rosenshine I, Finlay BB. In vitro and in vivo model systems for studying enteropathogenic Escherichia coli infections. Cold Spring Harb Perspect Med. 2013;3(3):a009977. Epub 2013/03/05. doi: 10.1101/cshperspect.a009977. PubMed PMID: 23457294; PubMed Central PMCID: PMC3579205.

21. Moon HW, Whipp SC, Argenzio RA, Levine MM, Giannella RA. Attaching and effacing activities of rabbit and human enteropathogenic Escherichia coli in pig and rabbit intestines. Infect Immun. 1983;41(3):1340–51. Epub 1983/09/01. PubMed PMID: 6350186; PubMed Central PMCID: PMC264644.

22. Larsen PL, Albert PS, Riddle DL. Genes that regulate both development and longevity in Caenorhabditis elegans. Genetics. 1995;139(4):1567–83. Epub 1995/04/01. PubMed PMID: 7789761; PubMed Central PMCID: PMC1206485.

23. Dean-Nystrom EA, Gansheroff LJ, Mills M, Moon HW, O’Brien AD. Vaccination of pregnant dams with intimin(O157) protects suckling piglets from Escherichia coli O157:H7 infection. Infect Immun. 2002;70(5):2414–8. Epub 2002/04/16. PubMed PMID: 11953378; PubMed Central PMCID: PMC127944.

24. Misyurina O, Asper DJ, Deng W, Finlay BB, Rogan D, Potter AA. The role of Tir, EspA, and NleB in the colonization of cattle by Shiga toxin producing Escherichia coli O26:H11. Can J Microbiol. 2010;56(9):739–47. Epub 2010/10/06. doi: 10.1139/w10-059. PubMed PMID: 20921984.

25. Savkovic SD, Villanueva J, Turner JR, Matkowskyj KA, Hecht G. Mouse model of enteropathogenic Escherichia coli infection. Infect Immun. 2005;73(2):1161–70. Epub 2005/01/25. doi: 10.1128/IAI.73.2.1161-1170.2005. PubMed PMID: 15664959; PubMed Central PMCID: PMC546940.

26. Royan SV, Jones RM, Koutsouris A, Roxas JL, Falzari K, Weflen AW, et al. Enteropathogenic E. coli non-LEE encoded effectors NleH1 and NleH2 attenuate NF-kappaB activation. Mol Microbiol. 2010;78(5):1232–45. Epub 2010/11/26. doi: 10.1111/j.1365-2958.2010.07400.x. PubMed PMID: 21091507; PubMed Central PMCID: PMC3325542.

27. Zhang Q, Li Q, Wang C, Liu X, Li N, Li J. Enteropathogenic Escherichia coli changes distribution of occludin and ZO-1 in tight junction membrane microdomains in vivo. Microbial pathogenesis. 2010;48(1):28–34. Epub 2009/10/17. doi: 10.1016/j.micpath.2009.10.002. PubMed PMID: 19833191.

28. Rhee KJ, Cheng H, Harris A, Morin C, Kaper JB, Hecht G. Determination of spatial and temporal colonization of enteropathogenic E. coli and enterohemorrhagic E. coli in mice using bioluminescent in vivo imaging. Gut Microbes. 2011;2(1):34–41. doi: 10.4161/gmic.2.1.14882. PubMed PMID: 21637016;PubMed Central PMCID: PMCPMC3225795.

29. Manthey CF, Autran CA, Eckmann L, Bode L. Human milk oligosaccharides protect against enteropathogenic Escherichia coli attachment in vitro and EPEC colonization in suckling mice. J Pediatr Gastroenterol Nutr. 2014;58(2):165–8. Epub 2013/09/21. doi: 10.1097/MPG.0000000000000172. PubMed PMID: 24048169.

30. Dupont A, Sommer F, Zhang K, Repnik U, Basic M, Bleich A, et al. Age-Dependent Susceptibility to Enteropathogenic Escherichia coli (EPEC) Infection in Mice. PLoS Pathog. 2016;12(5):e1005616. Epub 2016/05/10. doi: 10.1371/journal.ppat.1005616. PubMed PMID: 27159323; PubMed Central PMCID: PMC4861285.

31. Zhang Q, Li Q, Wang C, Li N, Li J. Redistribution of tight junction proteins during EPEC infection in vivo. Inflammation. 2012;35(1):23–32. Epub 2010/12/21. doi: 10.1007/s10753-010-9285-1. PubMed PMID: 21170673.

32. Bolick DT, Medeiros P, Ledwaba SE, Lima AAM, Nataro JP, Barry EM, et al. Critical Role of Zinc in a New Murine Model of Enterotoxigenic Escherichia coli Diarrhea. Infect Immun. 2018;86(7). Epub 2018/04/18. doi: 10.1128/IAI.00183-18. PubMed PMID: 29661930; PubMed Central PMCID: PMCPMC6013668.

33. Giallourou N, Medlock GL, Bolick DT, Medeiros PH, Ledwaba SE, Kolling GL, et al. A novel mouse model of Campylobacter jejuni enteropathy and diarrhea. PLoS Pathog. 2018;14(3):e1007083. Epub 2018/05/24. doi: 10.1371/journal.ppat.1007083. PubMed PMID: 29791507.

34. Ph QSM, Ledwaba SE, Bolick DT, Giallourou N, Yum LK, Costa DVS, et al. A murine model of diarrhea, growth impairment and metabolic disturbances with Shigella flexneri infection and the role of zinc deficiency. Gut Microbes. 2019:1–16. Epub 2019/02/05. doi: 10.1080/19490976.2018.1564430. PubMed PMID: 30712505.

35. Kosek MN, Investigators M-EN. Causal Pathways from Enteropathogens to Environmental Enteropathy: Findings from the MAL-ED Birth Cohort Study. EBioMedicine. 2017;18:109–17. Epub 2017/04/12. doi: 10.1016/j.ebiom.2017.02.024. PubMed PMID: 28396264; PubMed Central PMCID: PMC5405169.

36. McCormick BJ, Lee GO, Seidman JC, Haque R, Mondal D, Quetz J, et al. Dynamics and Trends in Fecal Biomarkers of Gut Function in Children from 1–24 Months in the MAL-ED Study. Am J Trop Med Hyg. 2017;96(2):465–72. Epub 2016/12/21. doi: 10.4269/ajtmh.16-0496. PubMed PMID: 27994110; PubMed Central PMCID: PMCPMC5303054.

37. Fahim SM, Das S, Gazi MA, Mahfuz M, Ahmed T. Association of intestinal pathogens with faecal markers of environmental enteric dysfunction among slum-dwelling children in the first 2 years of life in Bangladesh. Trop Med Int Health. 2018;23(11):1242–50. Epub 2018/08/23. doi: 10.1111/tmi.13141. PubMed PMID: 30133067; PubMed Central PMCID: PMCPMC6282798.

38. Iqbal NT, Sadiq K, Syed S, Akhund T, Umrani F, Ahmed S, et al. Promising Biomarkers of Environmental Enteric Dysfunction: A Prospective Cohort study in Pakistani Children. Sci Rep. 2018;8(1):2966. Epub 2018/02/16. doi: 10.1038/s41598-018-21319-8. PubMed PMID: 29445110; PubMed Central PMCID: PMCPMC5813024.

39. Aden K, Rehman A, Falk-Paulsen M, Secher T, Kuiper J, Tran F, et al. Epithelial IL-23R Signaling Licenses Protective IL-22 Responses in Intestinal Inflammation. Cell Rep. 2016;16(8):2208–18. Epub 2016/08/16. doi: 10.1016/j.celrep.2016.07.054. PubMed PMID: 27524624; PubMed Central PMCID: PMCPMC5443566.

40. Griesinger AM, Josephson RJ, Donson AM, Mulcahy Levy JM, Amani V, Birks DK, et al. Interleukin-6/STAT3 Pathway Signaling Drives an Inflammatory Phenotype in Group A Ependymoma. Cancer Immunol Res. 2015;3(10):1165–74. Epub 2015/05/15. doi: 10.1158/2326-6066.CIR-15-0061. PubMed PMID: 25968456; PubMed Central PMCID: PMCPMC4596749.

41. Westbom CM, Shukla A, MacPherson MB, Yasewicz EC, Miller JM, Beuschel SL, et al. CREB-induced inflammation is important for malignant mesothelioma growth. Am J Pathol. 2014;184(10):2816–27. Epub 2014/08/12. doi: 10.1016/j.ajpath.2014.06.008. PubMed PMID: 25111229; PubMed Central PMCID: PMCPMC4188864.

42. Zhang S, Xu W, Wang H, Cao M, Li M, Zhao J, et al. Inhibition of CREB-mediated ZO-1 and activation of NF-kappaB-induced IL-6 by colonic epithelial MCT4 destroys intestinal barrier function. Cell Prolif. 2019;52(6):e12673. Epub 2019/08/17. doi: 10.1111/cpr.12673. PubMed PMID: 31418947; PubMed Central PMCID: PMCPMC6869122.

43. Johansen K, Stintzing G, Magnusson KE, Sundqvist T, Jalil F, Murtaza A, et al. Intestinal permeability assessed with polyethylene glycols in children with diarrhea due to rotavirus and common bacterial pathogens in a developing community. J Pediatr Gastroenterol Nutr. 1989;9(3):307–13. Epub 1989/10/01. doi: 10.1097/00005176-198910000-00008. PubMed PMID: 2693681.

44. Rogawski EL J.; Platts-Mills, J.A.; Kabir, F.; Lertsethtakarn, P.; Siguas, M.; Khan, S.S.; Praharaj, I.; Murei, A.; Nshama, R.; et al. . Impact of enteropathogen infections on linear growth in children in low-resource settings using quantitative molecular diagnostics: Results from the MAL-ED cohort study. The Lancet Global Health. 2018;In Press.

45. Luperchio SA, Schauer DB. Molecular pathogenesis of Citrobacter rodentium and transmissible murine colonic hyperplasia. Microbes Infect. 2001;3(4):333–40. Epub 2001/05/04. doi: 10.1016/s1286-4579(01)01387-9. PubMed PMID: 11334751.

46. Mundy R, Girard F, FitzGerald AJ, Frankel G. Comparison of colonization dynamics and pathology of mice infected with enteropathogenic Escherichia coli, enterohaemorrhagic E. coli and Citrobacter rodentium. FEMS Microbiol Lett. 2006;265(1):126–32. Epub 2006/10/13. doi: 10.1111/j.1574-6968.2006.00481.x. PubMed PMID: 17034412.

47. Vulcano M, Dusi S, Lissandrini D, Badolato R, Mazzi P, Riboldi E, et al. Toll receptor-mediated regulation of NADPH oxidase in human dendritic cells. J Immunol. 2004;173(9):5749–56. Epub 2004/10/21. doi: 10.4049/jimmunol.173.9.5749. PubMed PMID: 15494527.

48. Bolick DT, Medeiros P, Ledwaba SE, Lima AAM, Nataro JP, Barry EM, et al. The Critical Role of Zinc in a New Murine Model of Enterotoxigenic E. coli (ETEC) Diarrhea. Infect Immun. 2018. Epub 2018/04/18. doi: 10.1128/IAI.00183-18. PubMed PMID: 29661930.

49. Guerrant RL, Leite AM, Pinkerton R, Medeiros PH, Cavalcante PA, DeBoer M, et al. Biomarkers of Environmental Enteropathy, Inflammation, Stunting, and Impaired Growth in Children in Northeast Brazil. PLoS One. 2016;11(9):e0158772. Epub 2016/10/01. doi: 10.1371/journal.pone.0158772. PubMed PMID: 27690129; PubMed Central PMCID: PMCPMC5045163.

50. Makhezer N, Ben Khemis M, Liu D, Khichane Y, Marzaioli V, Tlili A, et al. NOX1-derived ROS drive the expression of Lipocalin-2 in colonic epithelial cells in inflammatory conditions. Mucosal Immunol. 2019;12(1):117–31. Epub 2018/10/04. doi: 10.1038/s41385-018-0086-4. PubMed PMID: 30279516.

51. Dann SM, Spehlmann ME, Hammond DC, Iimura M, Hase K, Choi LJ, et al. IL-6-dependent mucosal protection prevents establishment of a microbial niche for attaching/effacing lesion-forming enteric bacterial pathogens. J Immunol. 2008;180(10):6816–26. Epub 2008/05/06. doi: 10.4049/jimmunol.180.10.6816. PubMed PMID: 18453602; PubMed Central PMCID: PMCPMC2696063.

52. Liu Z, Zaki MH, Vogel P, Gurung P, Finlay BB, Deng W, et al. Role of inflammasomes in host defense against Citrobacter rodentium infection. The Journal of biological chemistry. 2012;287(20):16955–64. Epub 2012/03/31. doi: 10.1074/jbc.M112.358705. PubMed PMID: 22461621; PubMed Central PMCID: PMCPMC3351318.

53. Sham HP, Yu EY, Gulen MF, Bhinder G, Stahl M, Chan JM, et al. SIGIRR, a negative regulator of TLR/IL-1R signalling promotes Microbiota dependent resistance to colonization by enteric bacterial pathogens. PLoS Pathog. 2013;9(8):e1003539. Epub 2013/08/21. doi: 10.1371/journal.ppat.1003539. PubMed PMID: 23950714; PubMed Central PMCID: PMCPMC3738496.

54. Gabay C, Lamacchia C, Palmer G. IL-1 pathways in inflammation and human diseases. Nat Rev Rheumatol. 2010;6(4):232–41. Epub 2010/02/24. doi: 10.1038/nrrheum.2010.4. PubMed PMID: 20177398.

55. Bolick DT, Whetzel AM, Skaflen M, Deem TL, Lee J, Hedrick CC. Absence of the G protein-coupled receptor G2A in mice promotes monocyte/endothelial interactions in aorta. Circ Res. 2007;100(4):572–80. Epub 2007/01/27. doi: 10.1161/01.RES.0000258877.57836.d2. PubMed PMID: 17255525.

56. Pezoldt J, Pisano F, Heine W, Pasztoi M, Rosenheinrich M, Nuss AM, et al. Impact of CCR7 on T-Cell Response and Susceptibility to Yersinia pseudotuberculosis Infection. J Infect Dis. 2017;216(6):752–60. Epub 2017/03/23. doi: 10.1093/infdis/jix037. PubMed PMID: 28329174.

57. Lee JH, Kim B, Jin WJ, Kim HH, Ha H, Lee ZH. Pathogenic roles of CXCL10 signaling through CXCR3 and TLR4 in macrophages and T cells: relevance for arthritis. Arthritis Res Ther. 2017;19(1):163. Epub 2017/07/21. doi: 10.1186/s13075-017-1353-6. PubMed PMID: 28724396; PubMed Central PMCID: PMCPMC5518115.

58. Green DS, Young HA, Valencia JC. Current prospects of type II interferon gamma signaling and autoimmunity. The Journal of biological chemistry. 2017;292(34):13925–33. Epub 2017/06/28. doi: 10.1074/jbc.R116.774745. PubMed PMID: 28652404; PubMed Central PMCID: PMCPMC5572907.

59. Long KZ, Rosado JL, Santos JI, Haas M, Al Mamun A, DuPont HL, et al. Associations between mucosal innate and adaptive immune responses and resolution of diarrheal pathogen infections. Infect Immun. 2010;78(3):1221–8. doi: 10.1128/IAI.00767-09. PubMed PMID: 20038536; PubMed Central PMCID: PMCPMC2825919.

60. Aychek T, Mildner A, Yona S, Kim KW, Lampl N, Reich-Zeliger S, et al. IL-23-mediated mononuclear phagocyte crosstalk protects mice from Citrobacter rodentium-induced colon immunopathology. Nature communications. 2015;6:6525. Epub 2015/03/13. doi: 10.1038/ncomms7525. PubMed PMID: 25761673; PubMed Central PMCID: PMCPMC4382688.

61. Wittkopf N, Pickert G, Billmeier U, Mahapatro M, Wirtz S, Martini E, et al. Activation of intestinal epithelial Stat3 orchestrates tissue defense during gastrointestinal infection. PLoS One. 2015;10(3):e0118401. Epub 2015/03/24. doi: 10.1371/journal.pone.0118401. PubMed PMID: 25799189; PubMed Central PMCID: PMCPMC4370566.

62. Alteri CJ, Mobley HL. Escherichia coli physiology and metabolism dictates adaptation to diverse host microenvironments. Curr Opin Microbiol. 2012;15(1):3–9. Epub 2011/12/30. doi: 10.1016/j.mib.2011.12.004. PubMed PMID: 22204808; PubMed Central PMCID: PMCPMC3265668.

63. Leonardi R, Zhang YM, Lykidis A, Rock CO, Jackowski S. Localization and regulation of mouse pantothenate kinase 2. FEBS letters. 2007;581(24):4639–44. Epub 2007/09/11. doi: 10.1016/j.febslet.2007.08.056. PubMed PMID: 17825826; PubMed Central PMCID: PMCPMC2034339.

64. Sugiharto S, Hedemann MS, Lauridsen C. Plasma metabolomic profiles and immune responses of piglets after weaning and challenge with E. coli. J Anim Sci Biotechnol. 2014;5(1):17. Epub 2014/03/15. doi: 10.1186/2049-18915-17. PubMed PMID: 24624922; PubMed Central PMCID: PMCPMC3995590.

65. Jeon SH, Lee MY, Rahman MM, Kim SJ, Kim GB, Park SY, et al. The antioxidant, taurine reduced lipopolysaccharide (LPS)-induced generation of ROS, and activation of MAPKs and Bax in cultured pneumocytes. Pulm Pharmacol Ther. 2009;22(6):562–6. Epub 2009/08/12. doi: 10.1016/j.pupt.2009.07.004. PubMed PMID: 19665057.

66. Oliveira MW, Minotto JB, de Oliveira MR, Zanotto-Filho A, Behr GA, Rocha RF, et al. Scavenging and antioxidant potential of physiological taurine concentrations against different reactive oxygen/nitrogen species. Pharmacol Rep. 2010;62(1):185–93. Epub 2010/04/03. doi: 10.1016/s1734-1140(10)70256-5. PubMed PMID: 20360629.

67. Swann JR, Tuohy KM, Lindfors P, Brown DT, Gibson GR, Wilson ID, et al. Variation in antibiotic-induced microbial recolonization impacts on the host metabolic phenotypes of rats. Journal of proteome research. 2011;10(8):3590–603. Epub 2011/05/20. doi: 10.1021/pr200243t. PubMed PMID: 21591676.

68. Kalliolias GD, Ivashkiv LB. TNF biology, pathogenic mechanisms and emerging therapeutic strategies. Nat Rev Rheumatol. 2016;12(1):49–62. Epub 2015/12/15. doi: 10.1038/nrrheum.2015.169. PubMed PMID: 26656660; PubMed Central PMCID: PMCPMC4809675.

69. Shifflett DE, Clayburgh DR, Koutsouris A, Turner JR, Hecht GA. Enteropathogenic E. coli disrupts tight junction barrier function and structure in vivo. Laboratory investigation; a journal of technical methods and pathology. 2005;85(10):1308–24. Epub 2005/08/30. doi: 10.1038/labinvest.3700330. PubMed PMID: 16127426.

70. Chelakkot C, Ghim J, Ryu SH. Mechanisms regulating intestinal barrier integrity and its pathological implications. Exp Mol Med. 2018;50(8):103. Epub 2018/08/18. doi: 10.1038/s12276-018-0126-x. PubMed PMID: 30115904; PubMed Central PMCID: PMCPMC6095905.

71. Smyth D, Phan V, Wang A, McKay DM. Interferon-gamma-induced increases in intestinal epithelial macromolecular permeability requires the Src kinase Fyn. Laboratory investigation; a journal of technical methods and pathology. 2011;91(5):764–77. Epub 2011/02/16. doi: 10.1038/labinvest.2010.208. PubMed PMID: 21321534.

72. Wang Y, Mumm JB, Herbst R, Kolbeck R, Wang Y. IL-22 Increases Permeability of Intestinal Epithelial Tight Junctions by Enhancing Claudin-2 Expression. J Immunol. 2017;199(9):3316–25. Epub 2017/09/25. doi: 10.4049/jimmunol.1700152. PubMed PMID: 28939759.

73. Guttman JA, Samji FN, Li Y, Vogl AW, Finlay BB. Evidence that tight junctions are disrupted due to intimate bacterial contact and not inflammation during attaching and effacing pathogen infection in vivo. Infect Immun. 2006;74(11):6075–84. Epub 2006/09/07. doi: 10.1128/IAI.00721-06. PubMed PMID: 16954399;PubMed Central PMCID: PMCPMC1695516.

74. Ugalde-Silva P, Gonzalez-Lugo O, Navarro-Garcia F. Tight Junction Disruption Induced by Type 3 Secretion System Effectors Injected by Enteropathogenic and Enterohemorrhagic Escherichia coli. Front Cell Infect Microbiol. 2016;6:87. Epub 2016/09/09. doi: 10.3389/fcimb.2016.00087. PubMed PMID: 27606286; PubMed Central PMCID: PMCPMC4995211.

75. Luissint AC, Parkos CA, Nusrat A. Inflammation and the Intestinal Barrier: Leukocyte-Epithelial Cell Interactions, Cell Junction Remodeling, and Mucosal Repair. Gastroenterology. 2016;151(4):616–32. Epub 2016/07/21. doi: 10.1053/j.gastro.2016.07.008. PubMed PMID: 27436072; PubMed Central PMCID: PMCPMC5317033.

76. Gauthier A, Puente JL, Finlay BB. Secretin of the enteropathogenic Escherichia coli type III secretion system requires components of the type III apparatus for assembly and localization. Infect Immun. 2003;71(6):3310–9. Epub 2003/05/23. doi: 10.1128/iai.71.6.3310-3319.2003. PubMed PMID: 12761113; PubMed Central PMCID: PMCPMC155723.

77. Zarivach R, Vuckovic M, Deng W, Finlay BB, Strynadka NC. Structural analysis of a prototypical ATPase from the type III secretion system. Nat Struct Mol Biol. 2007;14(2):131–7. Epub 2007/01/24. doi: 10.1038/nsmb1196. PubMed PMID: 17237797.

78. Levine MM, Nataro JP, Karch H, Baldini MM, Kaper JB, Black RE, et al. The diarrheal response of humans to some classic serotypes of enteropathogenic Escherichia coli is dependent on a plasmid encoding an enteroadhesiveness factor. The Journal of infectious diseases. 1985;152(3):550–9. Epub 1985/09/01. PubMed PMID: 2863318.

79. Jarvis KG, Giron JA, Jerse AE, McDaniel TK, Donnenberg MS, Kaper JB. Enteropathogenic Escherichia coli contains a putative type III secretion system necessary for the export of proteins involved in attaching and effacing lesion formation. Proc Natl Acad Sci U S A. 1995;92(17):7996–8000. Epub 1995/08/15. doi: 10.1073/pnas.92.17.7996. PubMed PMID: 7644527; PubMed Central PMCID: PMCPMC41273.

80. Zhang WL, Kohler B, Oswald E, Beutin L, Karch H, Morabito S, et al. Genetic diversity of intimin genes of attaching and effacing Escherichia coli strains. Journal of clinical microbiology. 2002;40(12):4486–92. Epub 2002/11/28. doi: 10.1128/jcm.40.12.4486-4492.2002. PubMed PMID: 12454140; PubMed Central PMCID: PMCPMC154638.

81. Livak KJ, Schmittgen TD. Analysis of relative gene expression data using real-time quantitative PCR and the 2(-Delta Delta C(T)) Method. Methods. 2001;25(4):402–8. Epub 2002/02/16. doi: 10.1006/meth.2001.1262. PubMed PMID: 11846609.

